# Pathogenic aggregates alter actin organization and cellular viscosity resulting in stalled clathrin mediated endocytosis

**DOI:** 10.1101/2023.07.10.548473

**Authors:** Surya Bansi Singh, Shatruhan Singh Rajput, Aditya Sharma, Vaishnavi Ananthanarayanan, Amitabha Nandi, Shivprasad Patil, Amitabha Majumdar, Deepa Subramanyam

## Abstract

Protein aggregation is a common underlying feature of neurodegenerative disorders. Cells expressing neurodegeneration–associated mutant proteins show altered uptake of ligands, suggestive of impaired endocytosis, in a manner as yet unknown. Using live cell imaging, we show that clathrin-mediated endocytosis (CME) is affected due to altered actin cytoskeletal organization in the presence of Huntingtin aggregates. Additionally, we find that cells containing Huntingtin aggregates are stiffer and less viscous than their wild-type counterparts due to altered actin conformation, and not merely due to the physical presence of aggregate(s). We further demonstrate that CME and cellular viscosity can be rescued by overexpressing Hip1, Arp2/3 or transient LatrunculinA treatment. Examination of other pathogenic aggregates revealed that only a subset of these display defective CME, along with altered actin organization and increased stiffness. Together, our results point to an intimate connection between functional CME, actin organization and cellular stiffness in the context of neurodegeneration.

## Introduction

Neurodegenerative diseases are conditions where the function of the nervous system is adversely affected due to progressive loss of neurons. Aggregation and misfolding of proteins is often linked to a number of neurodegenerative diseases such as Huntington’s disease, amyotrophic lateral sclerosis, Alzheimer’s disease and Parkinson’s disorder^1–5^.

Huntington’s disease, an inherited neurodegenerative disorder, is characterized by the formation of neuronal intracellular inclusions and loss of striatal projection neurons^6^. The Huntingtin protein, a 350kDa protein which is central to this disease, is a product of the Huntingtin gene which is ubiquitously expressed, and is conserved across a wide range of species^7, 8^. The underlying cause of this disease is due to an abnormal expansion of the CAG tract (responsible for encoding glutamine, Q) in Exon 1 of the Huntingtin gene^9^, which may promote an abnormal protein conformation, resulting in the formation of aggregates^10^. However the disease is manifested only when expansion of the CAG tract passes the pathological threshold of ∼35 – 40 repeats. Pathogenic Huntingtin aggregates are observed in the nucleus and cytoplasm of affected neurons^10^. In yeast, Huntingtin aggregation varied with the length of PolyQ expansion, suggesting that aggregation and pathogenicity depended on PolyQ expansion and could be regulated by the expression of chaperone proteins^11^. Huntingtin aggregates are capable of spreading from cell – to – cell in the *Drosophila* brain, resulting in a loss of the neuronal population. However, neuronal loss could be prevented by blocking endocytosis in the recipient neurons or synaptic exocytosis^12^. Wild type Huntingtin is known to specifically enhance the vesicular transport of Brain Derived Neurotrophic Factor (BDNF) along microtubule paths, with reduction in wild type Huntingtin attenuating trafficking in neuronal cells^13^. Internalization of transferrin, a cargo internalized via clathrin mediated endocytosis (CME) was drastically reduced in the presence of aggregated forms of Huntingtin and other pathogenic polyQ aggregates^14^.

Clathrin mediated endocytosis (CME) is a critical process involved in the cellular uptake of various molecules from the extracellular milieu and the plasma membrane. CME plays a critical role in synaptic vesicle trafficking^15^, with clathrin coated structures (CCSs) being central to this process^16^. Alterations in CME have been implicated in a variety of disorders such as Huntington’s disease, spinocerebellar ataxia type I and Amyotrophic lateral sclerosis. Previous reports demonstrate that protein aggregates formed by mutant forms of Huntingtin (HTT), Ataxin-1 (Atx1) and Superoxide dismutase-1 (SOD1) can inhibit CME and AMPA receptor recycling^14^. The polyQ containing Huntingtin exon 1 fragment is also known to interact with some of the proteins involved in CME such as Clathrin heavy chain, Huntingtin interacting protein 1 (Hip1), Adaptor protein complex 2 alpha 2 subunit and Dynamin1^17^. Further, uptake of transferrin, a ligand that undergoes endocytosis was reduced in the presence of Huntington aggregates, while mutations in genes involved in endocytosis enhanced Huntington-associated toxicity^18^.

Actin is known to play a central and critical role in the context of CME^19^. Alterations in the organization of actin are known to regulate the physical properties of cells^20^, which in turn are shown to impact the health and function of cells. A recent study, using Huntingtin knockout mice demonstrated that loss of Huntingtin resulted in hyperactivation of LIM kinase and stabilized the Actin cytoskeleton in a Cofilin-dependent manner. HTT knockout mice also showed an impairment of synaptic AMPAR (AMPA receptor) expression and trafficking, along with reduction in cortical neuronal spine length^21^. Synapse formation at the neuromuscular junction was disrupted and was accompanied by a reduction in lifespan in a *Drosophila* model of Huntington’s disease^22^.

Previous studies using cell lines and animal model systems have shown defects in the uptake of cargo through endocytic processes in neurodegenerative diseases such as Huntington’s disease, Amyotrophic lateral sclerosis and Parkinson’s disease. However, the molecular mechanism underlying compromised endocytosis and trafficking in neurodegeneration still remains largely unknown. Additionally, as CME is a dynamic process, live imaging data is critical to understand how it is affected by protein aggregation associated with neurodegeneration. In the current study we set out to study how CME and actin dynamics were affected in the presence of pathogenic protein aggregates and whether a common mechanism operated in all aggregate-induced neurodegenerative conditions. Our study reveals that the movement and directionality of clathrin coated structures (CCSs) are affected in the presence of pathogenic Huntingtin aggregates, accompanied by alterations in actin flow. We further demonstrate that pathogenic forms of Huntingtin disrupt actin reorganization by sequestering Arp2/3 and actin, with overexpression of either Arp3 or Hip1 partially rescuing the phenotype along with restoring directional CCS movement. Changes in the state of actin organization are tightly correlated with the stiffness of the cells^20, 23–25^. Using atomic force microscopy, we measured the rheological response of single cells containing Huntingtin aggregates. It revealed an increased cellular stiffness and reduced fluidity in an actin-dependent manner. Interestingly, overexpression of Hip1 or transient treatment with Latrunculin A restored cellular stiffness and fluidity to levels comparable to wild type cells, indicating that the physical presence of the aggregate alone does not contribute to alterations in cellular properties, but rather is dependent on reorganization of the actin cytoskeleton. Further, upon testing the effects of other pathogenic aggregates on CME and actin organization and dynamics, we found a strong correlation between disruption of CCS dynamics, actin flow and loss of fluidity with enhanced cell stiffness. Together our results indicate that Huntingtin aggregates remodel the cellular actin cytoskeleton in a manner rendering the cells stiffer, where it is unable to assist CCS movement. We further demonstrate that an active remodeling of the actin cytoskeleton can override some of the detrimental effects of the aggregates with respect to endocytosis.

## Results

### Clathrin mediated endocytosis and clathrin coated structure dynamics are compromised in the presence of pathogenic Huntingtin polyQ aggregates

A number of models have been used to study the detrimental effects of mutations in the Huntingtin protein rendering it pathogenic. Previous reports have demonstrated that *Drosophila melanogaster* expressing mutant forms of the Huntingtin gene display many of the phenotypes observed in affected human patients, including accumulation of protein aggregates in cells and reduced life span^22^. Other reports have demonstrated that cells containing Huntingtin aggregates display reduced transferrin uptake^18^ indicating that endocytosis may be compromised.

Clathrin mediated endocytosis (CME) is a highly dynamic process^26^. We used live cell imaging to better understand how movement of CCSs is affected in cells expressing HTTQ138. To track the movement of CCSs in real time, we used a transgenic *Drosophila* line in which clathrin light chain is fused to GFP and co-expressed this with pathogenic HTTQ138 or non-pathogenic HTTQ15. In the non-pathogenic HTTQ15 expressing hemocytes, we observed a centripetal movement of CCSs similar to previous observations^27^ (Fig. 1a, b). We found no difference in CCS movement between WT cells and cells expressing HTTQ15, and henceforth use these two cell types interchangeably (Fig. 1a and 2a and data not shown). However, in HTTQ138 expressing cells there was little to no movement of these CCSs (Fig 1a, c; Supplementary videos 1a and 1b). Using particle image velocimetry (PIV) analysis (see Materials and Methods for details and Supplementary Fig. 1b) to quantify the movement and directionality of CCSs, we determined that in HTTQ15 expressing cells, CCSs follow a specific path with defined speeds (Fig. 1b), whereas CCSs in HTTQ138 expressing cells do not show any directional movement, with negligible radial speed indicating a severe compromise in clathrin dynamics and movement of clathrin-coated vesicles (Fig 1c). Analysis of the distribution of flow-field directions obtained from PIV analysis relative to the polar direction revealed that the angles were sharply distributed around a value of 180°, showing the centripetal movement of CCSs (Fig. 1b, right). However, in the case of HTTQ138 hemocytes, a broad distribution of the angles was obtained, indicating the absence of any directional centripetal movement of CCSs (Fig. 1c, right). To check whether ligand uptake was also reduced, we determined the ability of hemocytes to internalize maleylated BSA (mBSA), a cargo for the anionic ligand binding scavenger receptor. HTTQ138 expressing hemocytes showed reduced colocalization of mBSA with CCSs coupled with reduced internalization in comparison to control hemocytes (Fig. 1d, e).

**Figure 1.**
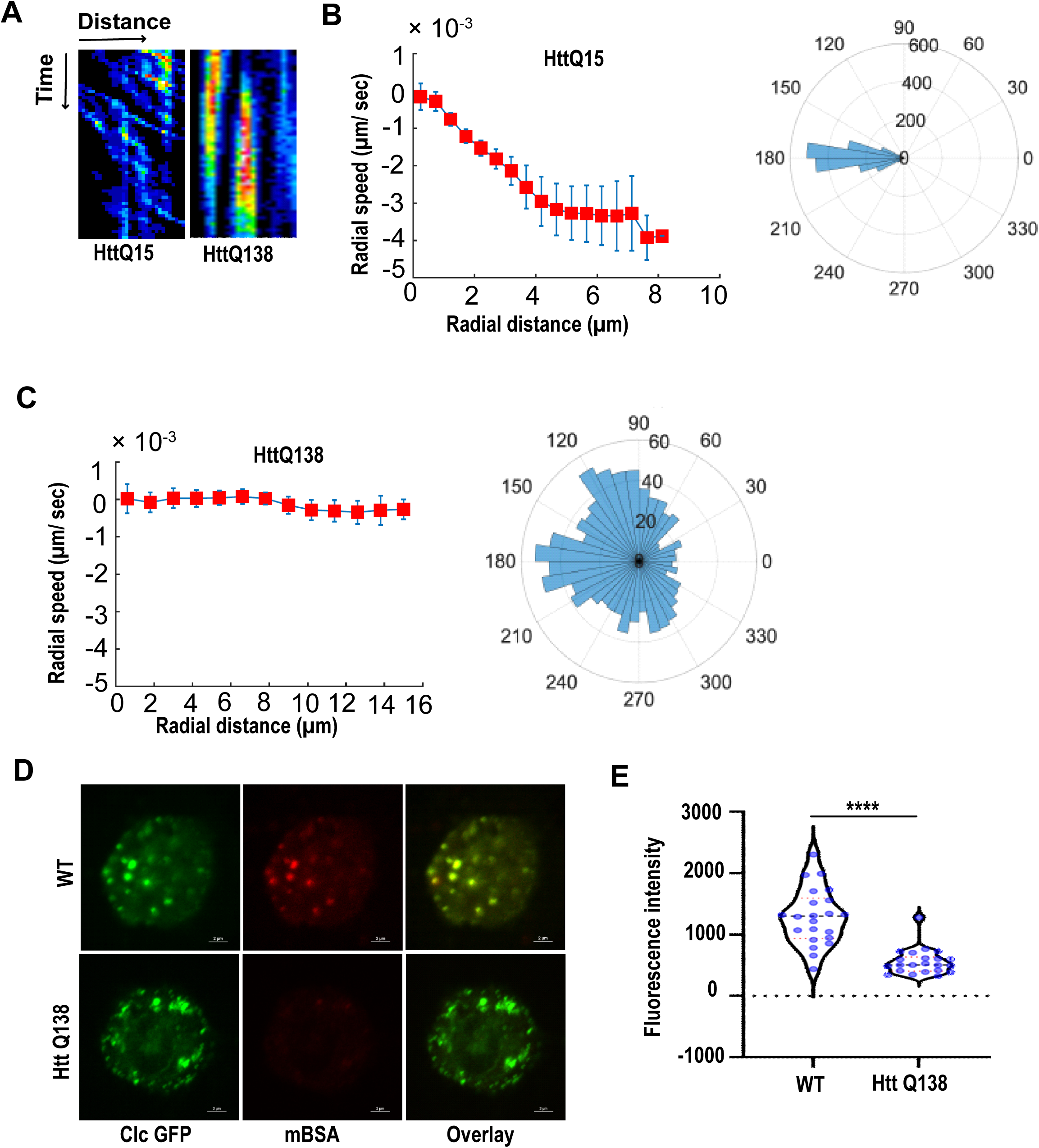
**A)** Kymograph showing movement of CCSs by live cell imaging of clathrin light chain tagged with GFP in HTTQ15 hemocytes and in presence of HTTQ138 aggregates. X-axis represents distance and Y-axis represents time. **B)** Radial speed (left) (µm/ sec) of the CCSs in HTTQ15 hemocytes as a function of radial distance (in µm) from the cell center obtained from time-averaged PIV analysis (see Material and Methods for details). Polar histogram of distribution of the flow-field directions obtained from PIV analysis relative to the polar direction (see Material and Methods for details) (right). The angles are sharply distributed around a value of 180°, showing the centripetal movement of CCSs. **C)** Graph (left) showing stalled movement of CCSs in HTTQ138 hemocytes. Polar histogram (right) of flow-field directions obtained similar to the HTTQ15 case in B) gives a broad distribution of the angles, indicating the absence of any directional centripetal movement of CCSs in presence of HTTQ138. **D)** Internalization of mBSA in WT and HTT Q138 cells. Clathrin puncta are shown in green and mBSA in the red channel. Scale bar 2 µm. **E)** Graph shows the quantification of internalized mBSA. Y-axis represents the intensity of mBSA internalized under different conditions. N=20 cells. **** P value < 0.0001 (Mann-Whitney test).

**Figure 2.**
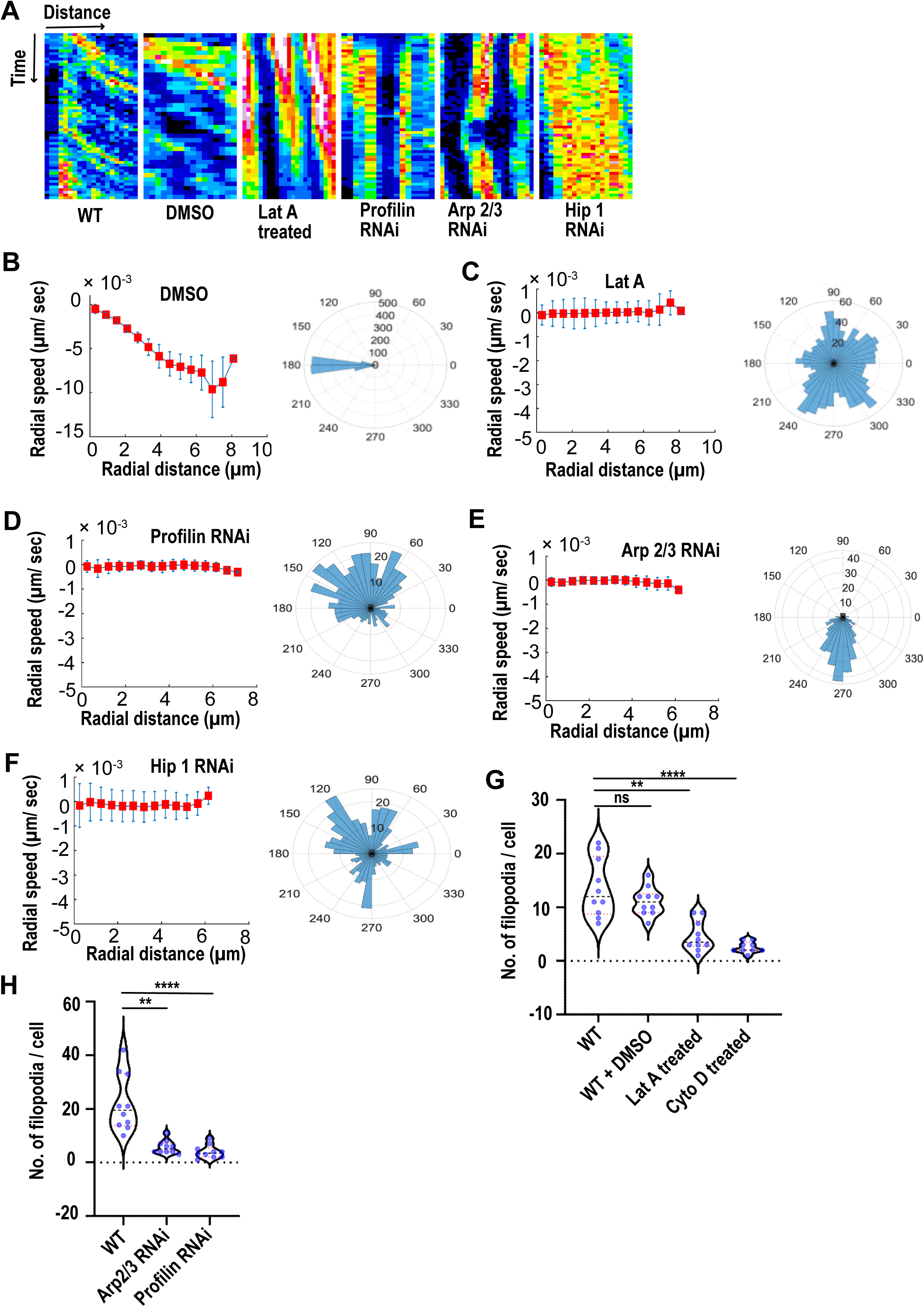
**A)** Kymographs showing movement of CCSs by live cell imaging of clathrin light chain tagged with GFP in WT, DMSO, Lat A treated, and on knockdown of Profilin, Arp2/3, Hip1. Radial speed (left) (µm/ sec) of the CCSs as a function of radial distance (in µm) from the cell center obtained from time-averaged PIV analysis similar to Fig. 1B, C in **B)** DMSO treated cells; **C)** LatA treated cells; **D)** Profilin knockdown cells; **E)** Arp2/3 knockdown cells; **F)** Hip1 knockdown cells. Polar histogram of distribution of the flow-field directions obtained from PIV analysis relative to the polar direction are shown to the right for each condition. The angles are sharply distributed around a value of 180°, showing the centripetal movement of CCSs in **B)**. Polar histogram of flow-field directions gives a broad distribution of the angles, indicating the absence of any directional centripetal movement of CCSs in **C-F**. **G)** Graph showing quantification of number of filopodia in WT, DMSO (vehicle control), LatA and CytoD treated cells. Number of cells = 10. ** P value = 0.0025, **** P value < 0.0001 (Kruskal-Wallis test followed by post hoc Dunn’s multiple comparisons test) **H)** Graph showing quantification of number of filopodia upon Arp2/3 and Profilin knockdown. Number of cells = 10. ** P value = 0.0025, **** P value < 0.0001 (Kruskal-Wallis test followed by post hoc Dunn’s multiple comparisons test.

### Huntingtin polyQ aggregates negatively affect Actin dynamics

We next addressed what might be a plausible explanation for the stagnation in CCS movement in HTTQ138 expressing cells. We hypothesized that this might be due to malfunction associated with cytoskeletal elements. Previously using inhibitors such as Latrunculin A, colchicine or nocodazole, it was shown that the centripetal CCS movement in hemocytes was dependent on the actin cytoskeleton and not on microtubules^27^. The role of the actin cytoskeleton is also well established in the context of CME, and CCS movement across various mammalian cell lines and in yeast^19, 28, 29^. Similar to what was previously reported, we found that hemocytes treated with pharmacological inhibitors of actin polymerization, Latrunculin A (LatA) or Cytochalasin D (CytoD) showed severe retardation in CCS movement in comparison to their DMSO treated counterparts (Fig. 2a and Supplementary Fig. 2a). This was also accompanied by a loss of CCSs speed and directional movement (Fig. 2b, c). As expected, actin dynamics in hemocytes treated with either Lat A or CytoD was also affected (data not shown). A reduction in the number of filopodia in Lat A and Cyto D treated cells (Fig. 2g) was also observed in comparison to DMSO treated cells.

To further understand how HTTQ138 impacted actin function, we coexpressed Actin-GFP along with HTTQ138. We observed sequestration of Actin-GFP in the HTT aggregates (Supplementary Fig. 1a). As this sequestration may result in removal and reduction of functional and available actin in the cells, we reasoned this might affect overall actin dynamics in HTTQ138 expressing cells. We performed live imaging of hemocytes coexpressing Lifeact-GFP along with either HTTQ15 or HTTQ138. While we observed a dynamic actin cytoskeleton accompanied by numerous filopodia extensions and retractions in WT or HTTQ15 expressing cells (Fig 3a, b, c, Supplementary video 2a and data not shown), there was a dramatic alteration in the dynamics of actin and reduced filopodia formation in HTTQ138 expressing cells (Fig. 3a, 3b, 3c Supplementary video 2b). PIV analysis of hemocytes expressing HTT Q138 aggregates also showed reduction in actin flow in comparison to wild type cells (Fig.3e).

**Figure 3.**
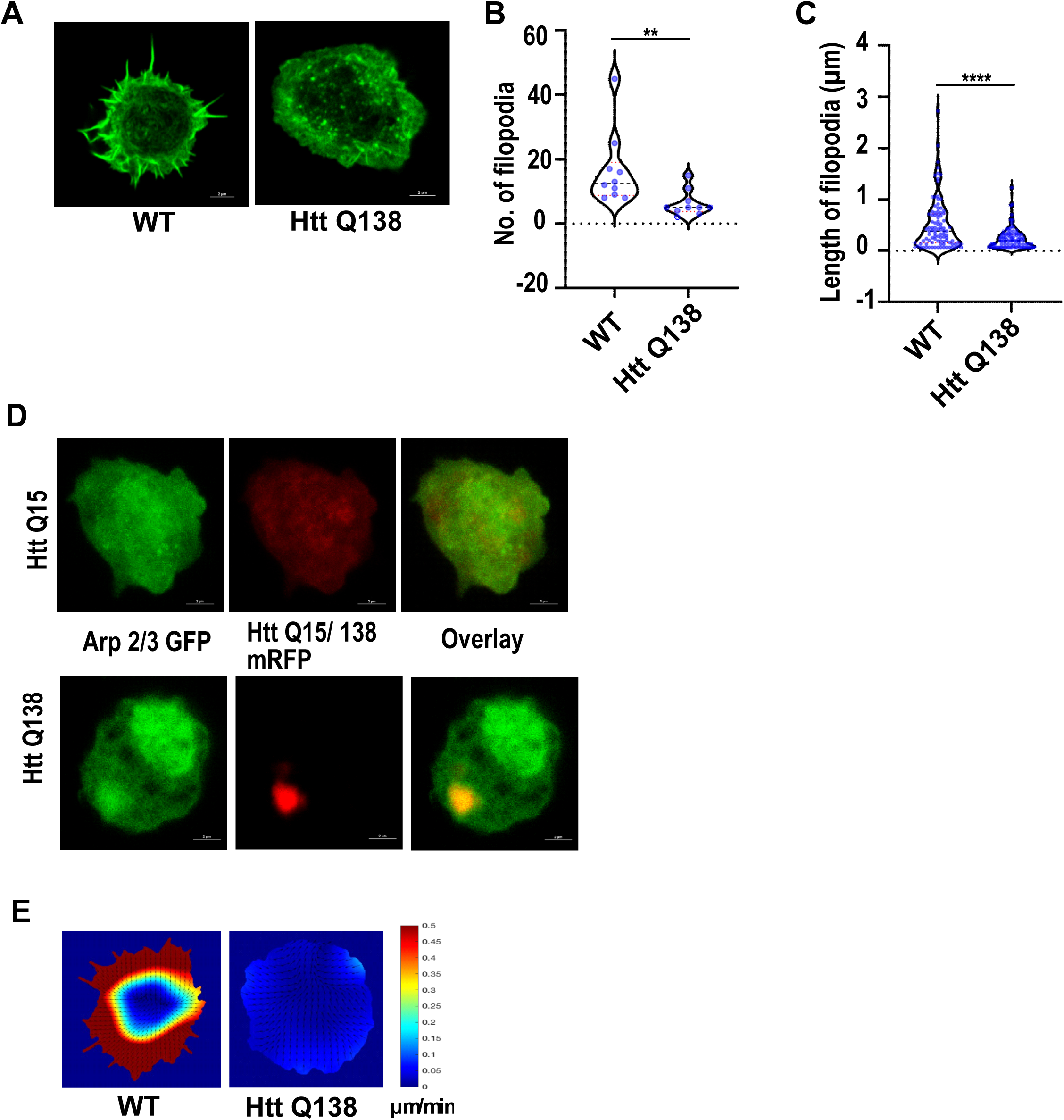
**A)** Representative micrograph showing filopodia formation in WT and HTT Q138 expressing hemocytes. **B, C)** Graphs showing the quantification of number and length of filopodia in WT and HTT Q138 expressing cells. Number of cells = 10, ** P value = 0.001 and **** P value < 0.0001 (Mann-Whitney test). **D)** Representative micrographs showing sequestration of Arp2/3 in HTT Q138 aggregates. Arp2/3 is shown in green and HTT Q15/ HTT Q138 - RFP is shown in red channel. **E)** Representative images showing actin flow in wild type and HTTQ138 expressing cells, obtained from PIV analysis.

Formation of filopodia requires the activity of the Arp2/3 complex^30^. The Arp2/3 complex consists of seven proteins namely, ArpC1, ArpC2, ArpC3, ArpC4, ArpC5, Arp2 and Arp3 and out of these Arp2 and Arp3 serve to nucleate actin, whereas the others bind to the existing actin filament and help in branching of the mother filament^31–33^. Since filopodia formation was compromised in HTTQ138 expressing cells, we decided to look at the distribution of the Arp2/3 complex by imaging hemocytes coexpressing Arp3-GFP along with HTTQ138. We found significant enrichment of Arp3 within HTT Q138 aggregates (Fig. 3d) indicating atleast a partial sequestration of the Arp2/3 complex. We reasoned if the sequestration of the Arp2/3 complex in HTTQ138 aggregates resulted in a loss of function of Arp2/3, and caused impaired CCS movement, we should see a similar impairment upon knocking down Arp3 directly in hemocytes even in the absence of HTTQ138. Upon inducing a transgenic Arp3 RNAi in hemocytes we observed a reduction in the number of filopodia (Fig 2h) along with stalled movement of CCSs (Fig. 2a) and loss of directional movement (Fig. 2e). Similar results were also obtained upon treatment of wild type cells with CK666, a chemical inhibitor of Arp2/3 (Supplementary Fig. 2a).

We also disrupted Actin polymerization genetically by targeting Profilin which is an Actin monomer binding protein and a key regulator of Actin polymerization^34, 35^. Knockdown of Profilin in hemocytes also resulted in an impairment in CCS movement as well as Actin dynamics (Fig. 2a, h). PIV analysis of CCSs upon Profilin knockdown also demonstrated defects in CCS movement and loss of their directionality (Fig. 2d). Chemical inhibition of Formin, a Rho GTPase involved in regulation of elongation of Actin fibers by SMIFH2, also resulted in stalled CCS movement (Supplementary Fig. 2a), further establishing that remodeling of the actin cytoskeleton is required for directional movement of CCSs.

Since, we found the impairment of CCS movement in HTTQ138 cells to be very similar to the phenotype that we observed on targeting the Actin polymerization pathways, we asked if the nature of individual CCSs in both these cases were also similar in terms of clathrin light chain exchange. Towards this we used fluorescence recovery after photo-bleaching (FRAP) to determine the dynamicity of Clathrin light chain exchange in the CCSs in the presence of pathogenic HTTQ138 and compared this to conditions where actin polymerization was perturbed. Recovery of fluorescence intensity of Clc-GFP was severely compromised in the presence of HTTQ138 (Supplementary Fig. 2b) compared to an almost complete recovery in wild type and control (Luc VAL 10) cells. Incomplete recovery was also observed upon knocking down Profilin or upon treatment with Lat A (Supplementary Fig. 2b). This indicates that targeting actin polymerization pathways can mimic the intracellular condition of the presence of pathogenic HTT aggregates by altering the exchange of clathrin light chain at CCSs.

Together, the observation of sequestration of Actin and Arp3 suggest a possible cytoplasmic reduction of Actin and Actin nucleating proteins as a likely cause for dysfunctional Actin dynamics and impairment of CCS movement in the presence of HTTQ138.

We also examined the involvement of microtubules in the context of HTTQ138. While microtubules are not generally considered to play a role in the early steps of Clathrin mediated endocytosis and clathrin coated vesicle dynamics^27^, other studies have demonstrated their involvement in receptor-mediated endocytosis^36–38^. We found that the organization and dynamics of microtubules remained unchanged even in the presence of Huntingtin aggregates (Supplementary video 2c and 2d) supporting the previous finding^27^ that microtubules do not play a role in early events of Clathrin mediated endocytosis.

We further speculated that the CCS movement on Actin cytoskeleton may involve specific motor proteins. Huntingtin is known to interact with Myosin VI via its optineurin binding domain^39^. The actin-based cytoskeleton motor, Myosin VI, is also implicated in various steps of membrane trafficking^40^ and shown to be involved in trafficking of clathrin coated vesicles^41^. The co-localization of myosin VI with actin polymerization regulatory proteins such as Cortactin and the Arp2/3 complex has also been shown suggesting a role for this motor in the regulation of actin dynamics. It has also been shown that Myosin VI is associated with CCSs through its C – terminal tail^42^. In Purkinje neurons, Myosin VI is known to promote Clathrin mediated endocytosis of AMPA receptors^43^. The coordinated action of myosin VI and Clathrin light chain a (CLCa) are essential for the fission of clathrin coated pits^44^. Knocking down Myosin VI, resulted in an impairment of CCS movement (Supplementary Fig. 2a and Supplementary movie 3a). However there was no discernible effect on actin dynamics or on filopodia formation (Supplementary movie 3b).

### Overexpression of Arp3, Hip1 and Mrj partially rescue CCSs movement in Huntingtin polyQ expressing cells

In order to determine whether the defective CCS movement observed in the presence of pathogenic HTT aggregates was predominantly driven by an alteration in actin dynamics due to sequestration of actin binding proteins by HTT aggregates, we asked whether overexpression of actin binding proteins could rescue the directional movement of CCSs. Towards this we coexpressed Arp3 along with pathogenic HTTQ138 in hemocytes and used live imaging to check the status of CCS movement and Actin dynamics in these cells. We observed a partial restoration of CCS movement and directionality along with filopodia formation (Fig. 4a, 4c, 4f, supplementary movie 4a and 4b).

**Figure 4.**
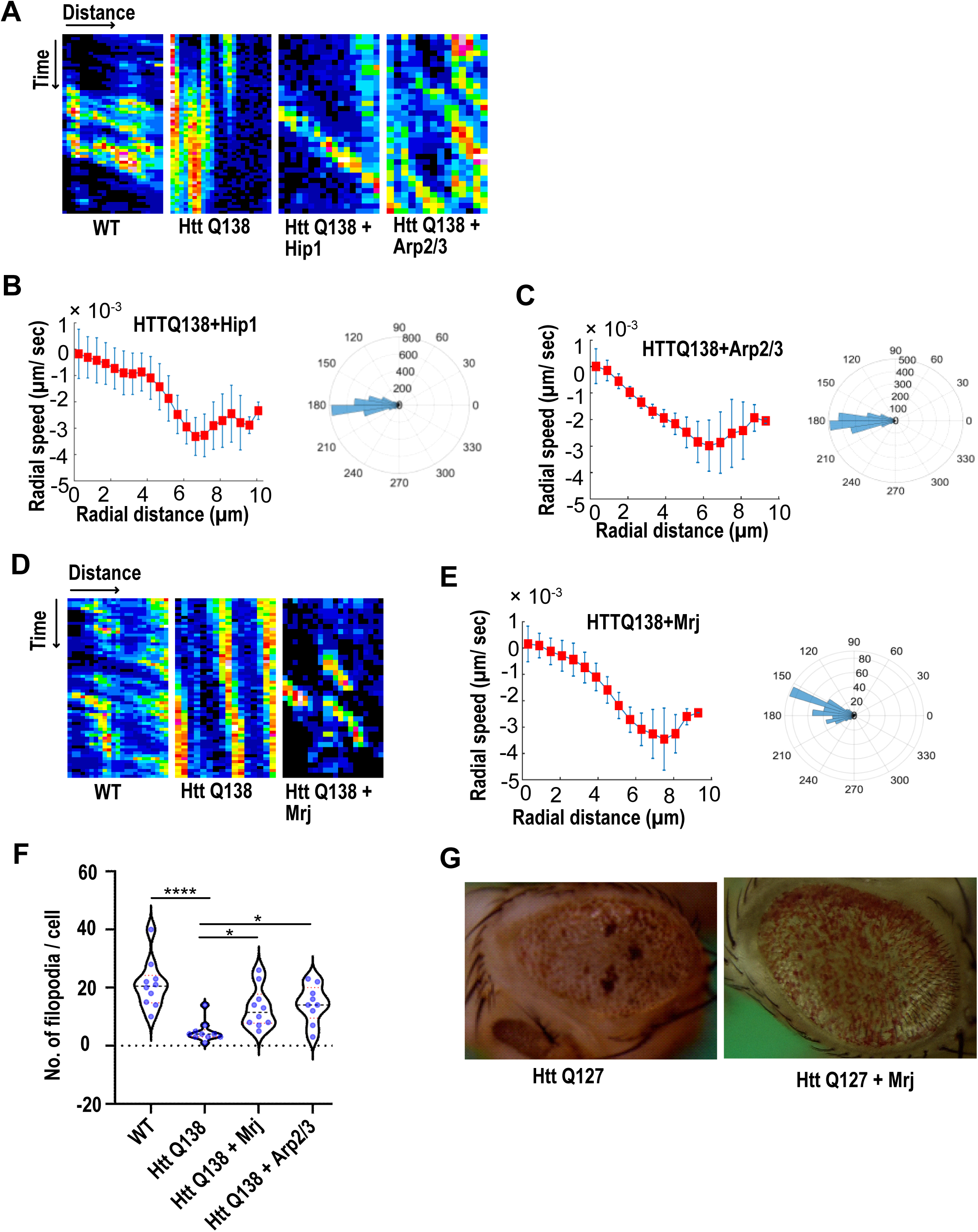
**A)** Kymographs showing movement of CCSs by live cell imaging of clathrin light chain tagged with GFP in WT, HTT Q138, HTT Q138 + Hip1 and HTT Q138 + Arp2/3 cells. Radial speed (left) (µm/ sec) of the CCSs as a function of radial distance (in µm) from the cell center obtained from time-averaged PIV analysis similar to Fig. 1B, C in **B)** HTT Q138 + Hip1; and **C)** HTTQ138 + Arp2/3 cells. Polar histogram of distribution of the flow-field directions obtained from PIV analysis relative to the polar direction are shown to the right for each condition. The angles are sharply distributed around a value of 180°, showing the centripetal movement of CCSs. **D)** Kymographs showing movement of CCSs by live cell imaging of clathrin light chain tagged with GFP in WT, HTT Q138, HTT Q138 + Mrj cells. **E)** Radial speed (left) (µm/ sec) of the CCSs as a function of radial distance (in µm) from the cell center obtained from time-averaged PIV analysis similar to Fig. 1B, C in HTTQ138 + Mrj cells. Polar histogram of distribution of the flow-field directions obtained from PIV analysis relative to the polar direction are shown to the right for each condition. The angles are sharply distributed around a value of 180°, showing the centripetal movement of CCSs. **F)** Box plot showing quantification of number of filopodia in WT, HTT Q138, HTT Q138 + Mrj and HTT Q138 + Arp2/3 cells. Number of cells = 10, * P value = 0.0309 for HTTQ138+Mrj and * P value = 0.0154 for HTTQ138+Arp2/3, **** P value < 0.0001 (Kruskal-Wallis test followed by post hoc Dunn’s multiple comparisons test). **G)** Representative micrograph showing *Drosophila* eye expressing HTTQ127 and HTTQ127+Mrj.

Huntingtin interacting protein1 (Hip1) is one of several proteins involved in the initial stages of Clathrin mediated endocytosis^45, 46^. Hip1 is also a known interactor of Huntingtin protein and expansions in the polyglutamate tract, as seen in pathological conditions, are known to disrupt this interaction^47^. Additionally, Hip1 is also known to interact with actin^48^ and the Hip1-actin interaction is known to be regulated by clathrin light chain ^49^. These studies indicate that Hip1 interaction with actin and clathrin is necessary for CME to occur under specific conditions. Upon coexpressing Hip1 with pathogenic HTTQ138, we observed a partial rescue of CCS movement (Fig. 4a, 4b, supplementary movie 4c). Conversely, knocking down Hip1 in wild type hemocytes resulted in a reduction in movement (Fig. 2a), speed, and loss of directionality of CCSs (Fig 2f), thus highlighting the importance of the Hip1-Actin axis in CME and its possible disruption in the presence of HTTQ138.

HTTQ138 forms aggregates inside the cell due to misfolding of protein, and its toxicity and aggregation was reported to be modulated by DNAJ chaperones^50, 51^. Previously, Drosophila Mrj, a homologue of mammalian DnaJB6 was observed to rescue polyglutamine associated toxicity in Drosophila. In a screen for polyQ modifiers, DNAJB6, a member of the DNAJ (HSP40) chaperone family was found to efficiently suppress polyQ aggregation in cells^52^. Brain-specific co-expression of DNAJB6 in a mouse model of Huntington’s disease delayed symptoms and increased the lifespan of Huntington mice^53^. Previous studies in mice and Drosophila have also shown that polyglutamine toxicity and aggregation can be modulated by DNAJ chaperones^50, 51^. The DNAJ-like protein Mrj, which is highly enriched in the brain, can effectively suppress polyQ toxicity^54^. Human DNAJB6 was also identified as an interactor of clathrin light chain a (CLTA) in a screen designed to determine the interactome of J domain-containing proteins^55^. Additionally, Mrj was also identified in an independent screen to modulate the cytotoxicity in the context of protein aggregation^56^. We found that coexpressing Mrj along with HTTQ138 in hemocytes partially restored CCS movement (Fig. 4d) speed and directionality (Fig. 4e, supplementary movie 4d) along with filopodia formation (Fig 4f). Overexpression of Mrj could also rescue the neurodegeneration observed in *Drosophila* eyes caused by the overexpression of pathogenic HTTQ127 (Fig. 4g). Together these results indicate that increasing the availability of proteins involved in actin reorganization are capable of restoring CME even in the presence of pathogenic aggregates.

### HTTQ138 expressing cells are stiffer with significantly reduced fluidity

A handful of previous studies have focused on studying the physical properties and transition of soluble oligomers into insoluble fibrils. In the past, atomic force microscopy (AFM) has been used to measure the kinetics of aggregate formation^57^. However, these have been carried out using purified proteins, and how these aggregates affect the physical state of a cell in which they are present, remains an unexplored question. Using AFM, we performed nanoindentation on wild type and HTT Q138 expressing cells to measure their stiffness and viscoelastic response. The experiments were performed on single cells using a spherical indenter - a glass-bead of 5µm diameter attached to the end of the tipless cantilever. The bead-attached cantilever was approached towards a single cell with a constant speed of 2 µm/s. After contact with the cell, the cantilever was pressed further to deform the cells by ∼ 500 nm and retracted back with the same speed. In the complete approach-retract cycle, the cantilever bending was recorded at every point of cantilever displacement which was then converted to force vs cell–deformation curve known as the force-curve. A representative force-curve on a wild type cell is shown in Supplementary Fig. 3a. Here, the points (a), (b), (c), (d), and (e) represent the conditions at various phases of cell deformations while the bead indents on the cell, with the bead leaving the cell surface at point (e), compared to the point at which it makes contact while approaching (a). Supplementary Figure 5a shows representative curves from each cell type. This represents slow recovery of cell shape and the hysteresis is indicator of the cell’s viscous (dissipative) response. We analyzed the force-curves by fitting them to Ting’s model, assuming that the cells follow a power-law-rheology (PLR) behavior (discussed in the Materials and Methods section). The basis for the PLR model is provided by the Soft glassy rheology (SGR) theory, which was developed to predict the behavior of soft glassy materials^58^. It assumes that the observed general scale-free behavior is a natural consequence of disorder and metastability of a material’s internal structure. Cytoskeleton, which consists of many disordered elements and are held together by weak attractive forces, may represent such a structure in cells^59^. Rheological behavior of cells is mediated by changes in the level of internal disorder and the effective temperature associated with cytoskeleton remodeling as suggested by SGR theory^60^. Hence, we surmised that actin dynamics and the rheological responses of cells are correlated and decided to perform rheological measurements on all cell types. The fitting procedure developed by Efremov et al^61^, for measurement of a cell’s rheology using AFM, returns two parameters-E_0_ and α, where E_0_ is a measure of the cell’s instantaneous modulus and α, the exponent in power law rheology. The exponent is related to the effective temperature of the material (amount of agitation energy in the system). Materials exhibit a solid-like behavior at α = 0, and a fluid-like behavior at α = 1. In other words, α can be interpreted as a measure of a cell’s fluidity. The availability of freely-moving cytoplasmic elements determines fluidity of a cell, for instance cells with freely moving actin elements are expected to be more liquid-like compared to cells in which actin is present in the same amount but whose movement is restricted.

Our results demonstrated that cells expressing HTTQ138 were stiffer than WT cells indicated by the higher Young’s modulus, E_0_ (Fig. 5a). The stiffness of cells expressing pathogenic Huntingtin aggregates was partially rescued upon overexpression of either Hip1 or Mrj (Fig. 5a) indicating that partial restoration in actin organization may also rescue stiffness of these cells in addition to CME. We also noticed a decreased stiffness of cells upon Lat A treatment in comparison to DMSO treated cells (Fig. 5a) suggesting that an altered state of the cytoskeleton was largely responsible for contributing to the increased stiffness observed in the presence of HTTQ138. We observed a correlation between α and E_0_. The enhanced E_0_ is accompanied by a reduction in α, indicating reduced fluidity (Fig. 5a, b). Fig. 5a shows stiffness variations in cells due to the presence of HTTQ138 aggregates and upon overexpression of other proteins mentioned in the previous section. These results demonstrate that depolymerization of the Actin cytoskeleton can reduce the cellular stiffness observed in the presence of HTTQ138. Our results suggest that due to the increased stiffness of HTTQ138 cells CCS movement may be impaired. Conversely, cells treated with LatA, in which the cytoskeletal architecture is completely broken down, are too soft and are hence also unable to support CCS movement. We therefore hypothesized that a brief, transient treatment of Lat A to HTT Q138 cells may result in a partial movement of CCSs at an intermediate time point where the stiff actin cytoskeleton is being converted to a more dynamic form. We were able to see movement of CCSs in these cells at an intermediate time point (Fig. 5c) thereby strengthening the idea that CCS movement requires a dynamic form of the actin cytoskeleton and that HTTQ138 causes the cytoskeleton to be held in a very stiff state. It should be noted here that the enhanced stiffness is that of a glassy state. The loss of fluidity accompanying the stiffness enhancement is an indicator of increased disorder with lower effective noise temperature. It can be said that the cells in general behave like a soft glass. The presence of aggregates lowers the effective temperature pushing it nearer to the glass transition, affecting transport.

**Figure 5.**
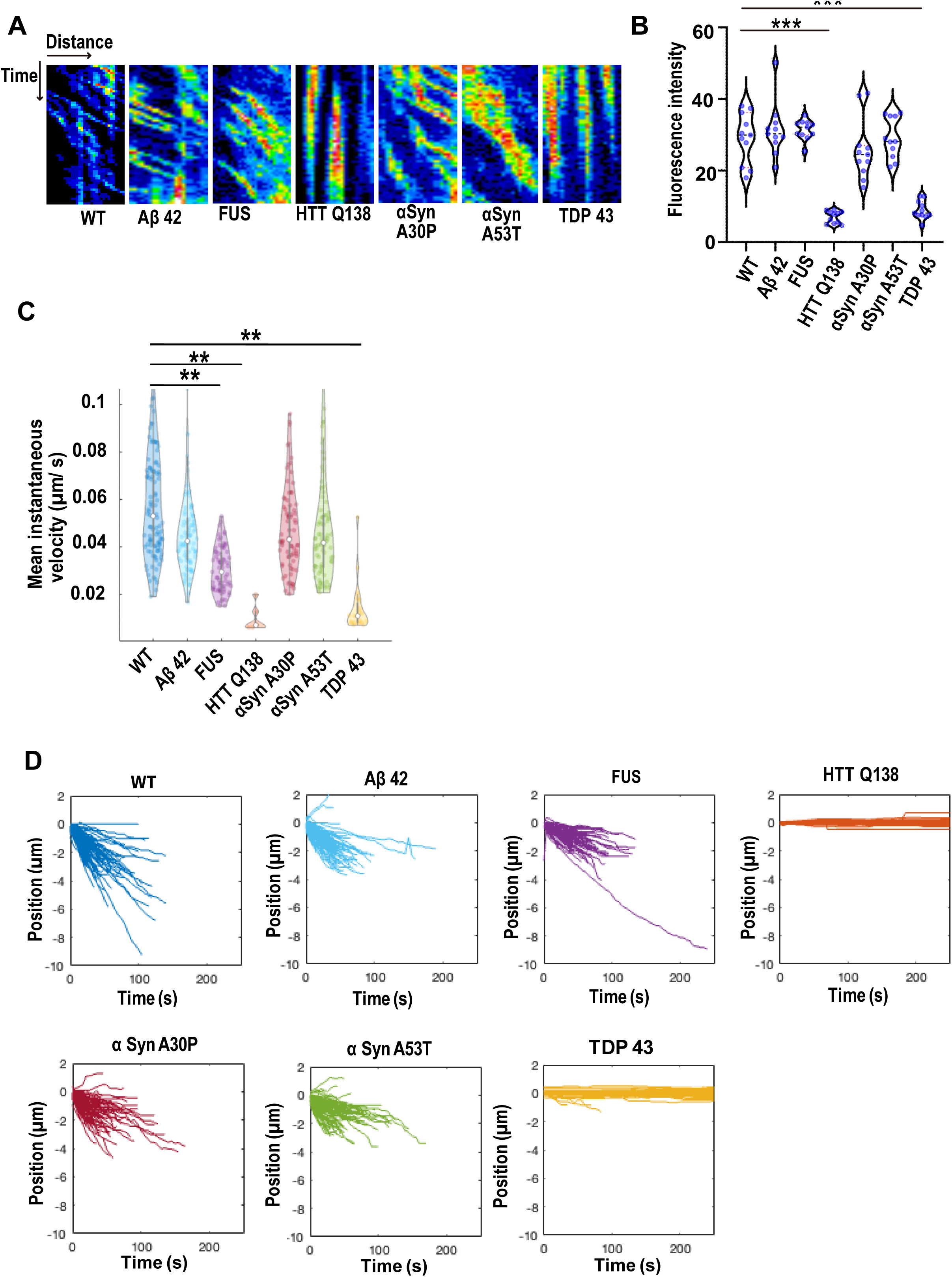
**A)** Box plot showing Young’s modulus of WT, DMSO, WT + Lat A, HTT Q138, HTT Q138 + Lat A, HTT Q138 + Hip1 and HTT Q138 + Mrj cells. Number of cells: WT = 9; WT+DMSO = 5; HTTQ138 = 10; WT+Lat A =10; HTTQ138+Lat A = 9; HTTQ138+Mrj = 6; HTTQ138+Hip1 = 7. * P value < 0.05, *** P value = 0.0007, ****, P value <0.0001 (Kruskal-Wallis test followed by post hoc Dunn’s multiple comparisons test). **B)** Graph shows the α value for wild type cells and cells treated as indicated or expressing specific combinations of proteins in the presence of HTTQ138. Number of cells: WT = 9 cells, WT+DMSO = 5 cells, HTTQ138 = 10 cells, WT+Lat A =10 cells, HTTQ138+Lat A = 9 cells, HTTQ138+Mrj = 6 cells, HTTQ138+Hip1 = 7 cells. * P value < 0.05 (Kruskal-Wallis test followed by post hoc Dunn’s multiple comparisons test). **C)** Kymographs showing movement of CCSs by live cell imaging of clathrin light chain tagged with GFP under indicated conditions.

### CCS movement and Actin dynamics is compromised in the presence of pathogenic TDP-43 similar to HTTQ138

As the presence of misfolded protein aggregates is also a hallmark of several other neurodegenerative conditions, we performed a systematic analysis of CCS movement and Actin dynamics in the presence of other such proteins. Towards this we coexpressed Aβ-42, FUSR521C, αSynA30P, αSynA53T, TDP-43 along with Clc-GFP and used live imaging to study CCS dynamics. We observed the while in the presence of Aβ-42, FUSR521C, αSynA30P or αSynA53T (Fig. 6a, Supplementary movies 5a, 5b, 5c, 5d) there was no defect in CCS movement, in the presence of TDP-43 we noticed a significant reduction in CCS movement (Fig.6a, 6d, Supplementary movie 5e). Tracking and quantification of CCSs in WT cells and cells expressing mutant proteins revealed that CCSs moved inwards in all cases, except TDP-43 expressing cells, in which movement was completely stalled (Fig. 6a, Supplementary movie 5e), similar to what we observed in the presence of HTTQ138. This was also supported by analysis of the mean instantaneous velocities, which showed a significant decrease in the presence of TDP-43 (Fig. 6c). Additionally, the uptake of fluorescently tagged mBSA was markedly reduced in the presence of TDP-43 (Fig.6b) indicative of a reduction in CME. In order to determine whether the recruitment or exchange of clathrin to CCSs was altered in the presence of aggregating pathogenic proteins, we performed FRAP on individual CCSs in hemocytes. Recovery of clathrin was severely compromised only in the presence of TDP-43, similar to HTTQ138, and was indistinguishable from WT in the context of other aggregates (Supplementary Fig. 4).

**Figure 6.**
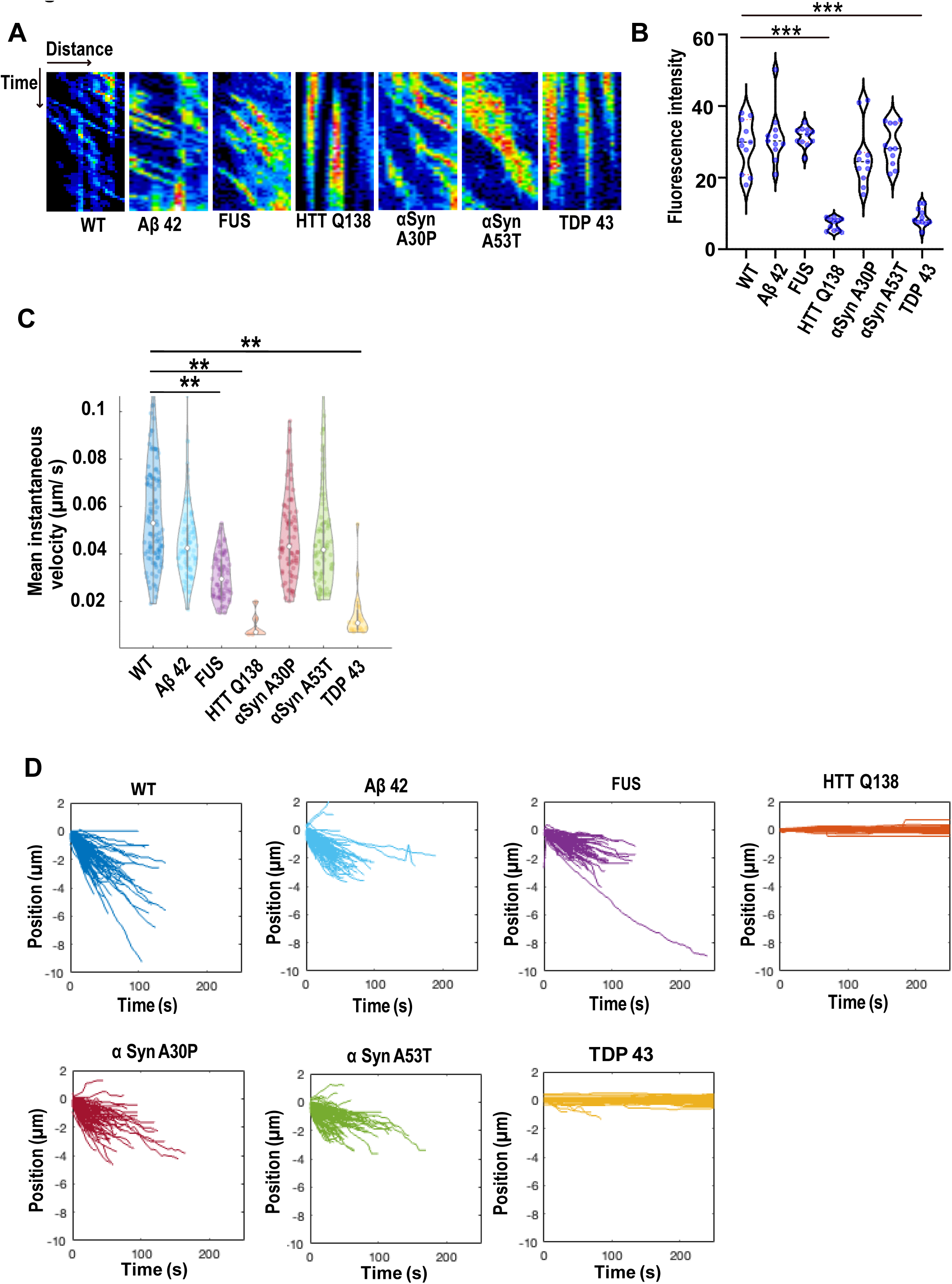
**A**) Kymographs showing the movement of CCSs from wild type hemocytes, or in the presence of the indicated proteins. **B)** Quantification of fluorescence intensity of internalized mBSA in hemocytes expressing the indicated aggregating proteins n=10). Statistical significance was determined using Kruskal-Wallis test followed by post hoc Dunn’s multiple comparison test. For WT vs HTT Q138 P value = 0.0001, *** and for WT vs TDP 43 P value = 0.0006, ***. **C)** Violin plots of mean instantaneous velocities of CCSs in WT hemocytes and hemocytes expressing aggregating proteins. Asterisks represent a significant difference from WT (P<0.05, Kruskal Wallis test for non-parametric data). **D)** Position vs. time plots of CCSs in WT hemocytes and hemocytes expressing aggregating proteins. The positions of CCSs with respect to the cell centre are plotted. Negative positions indicate movement towards the cell centre. The total number of vesicles used for quantitation from ten cells are as follows: WT – 117; Aβ42 – 105; FUSR521C – 83; HTTQ138 – 103; αSynA30P – 91; αSynA53T - 95 and TDP 43 – 106.

We also looked at Actin dynamics using LifeAct-GFP in the presence of these pathogenic proteins. Here again, we saw no significant difference in filopodia formation in the hemocytes expressing Aβ–42 (Fig. 7a, Supplementary movie 6a), FUSR521C (Fig. 7a, Supplementary movie 6b) or both mutants of α-Synuclein (Fig. 7a, Supplementary movie 6c and 6d), but noticed a dramatic reduction in the presence of TDP–43 (Fig. 7a, Supplementary movie 6e). Number, length and lifetime of filopodia were also compromised in the presence of TDP-43 expressing cells as compared to wild type cells (Fig. 7b, 7c, 7d), while remaining largely unaffected in the context of other mutants. Reduced actin flow was also observed in TDP-43 expressing cells in comparison to WT cells and cells expressing other aggregates (Fig. 7e). Together, these results indicate that CCS movement, CME and Actin dynamics are altered specifically in the presence of pathogenic TDP-43 and is phenotypically similar to our observations with HTTQ138.

**Figure 7.**
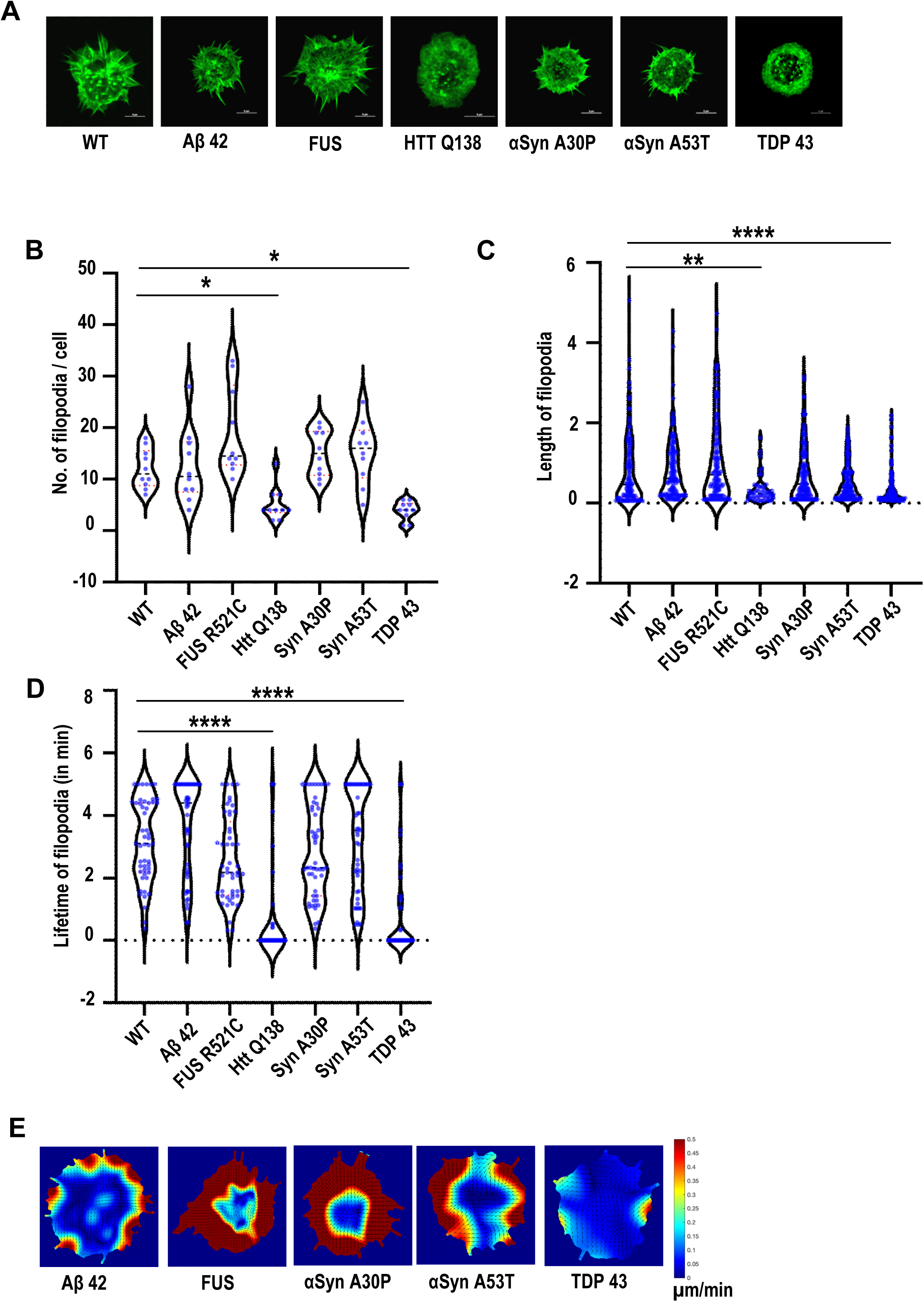
**A)** Representative micrographs showing the organization of actin marked by Lifeact-GFP in hemocytes. Scale bar 5µm in length. **B)** Graph showing the quantification of the number of filopodia per cell (n = 10 cells for each condition). Statistical significance was determined using Kruskal-Wallis test followed by post hoc Dunn’s multiple comparison test. For WT vs HTT Q138 P value = 0.0448,* and for WT vs TDP43 P value = 0.0135, *. **C)** Graph shows the quantification of length of filopodia in micrometres (n =10 cells for each condition). Statistical significance was determined using Kruskal-Wallis test followed by post hoc Dunn’s multiple comparison test. For WT vs HTT Q138 P value = 0.0041, ** and for WT vs TDP 43 P value < 0.0001, ****. The number and length of filopodia were quantified using the Filoquant plugin in ImageJ. **D)** Violin plots of the lifetime of filopodia in cells expressing Aβ42, FUSR521C, αSynA30P, αSynA53T, HTTQ138, TDP-43 compared to WT cells. Kruskal-Wallis test followed by post hoc Dunn’s multiple comparison was performed to calculate the P value. (**** P value < 0.0001). **E)** PIV analysis performed on LifeAct-GFP-expressing cells in the presence of the indicated pathogenic aggregating proteins to highlight the direction and magnitude of actin flow.

### TDP-43 aggregates result in the alteration of cellular physical properties

To elucidate the effect of cell’s pathogenic aggregates on its rheological response, we performed AFM experiments on single cells harboring different types of aggregates-­-Aβ–42, FUS R521C, αSynA30P, αSynA53T or TDP-43, similar to what was described in Fig. 5. Our PIV analysis has shown that, with the exception of HTTQ138 and TDP-43, the actin dynamics does not get altered in presence of other aggregates (Fig. 7). Both CME and filopodia formation were significantly altered in the presence of HTTQ138 and TDP-43, while Aβ–42, FUS R521C, αSynA30P and αSynA53T had no significant effects on CME or filopodia formation as seen in wild-type cells (Fig. 6 and 7). The AFM experiments showed a direct correlation of the cell’s physical properties with the state of actin organization and CME. E_0_ and α for cells having pathogenic aggregates of Aβ–42, FUS R521C, α-SynA30P and α-SynA53T remained similar to WT cells (Fig 8a,b). These parameters were altered in case of TDP-43, similar to what was seen in HTTQ138, showing a direct correlation of cell’s modulus of elasticity and fluidity with actin dynamics and CME transport.

**Figure 8.**
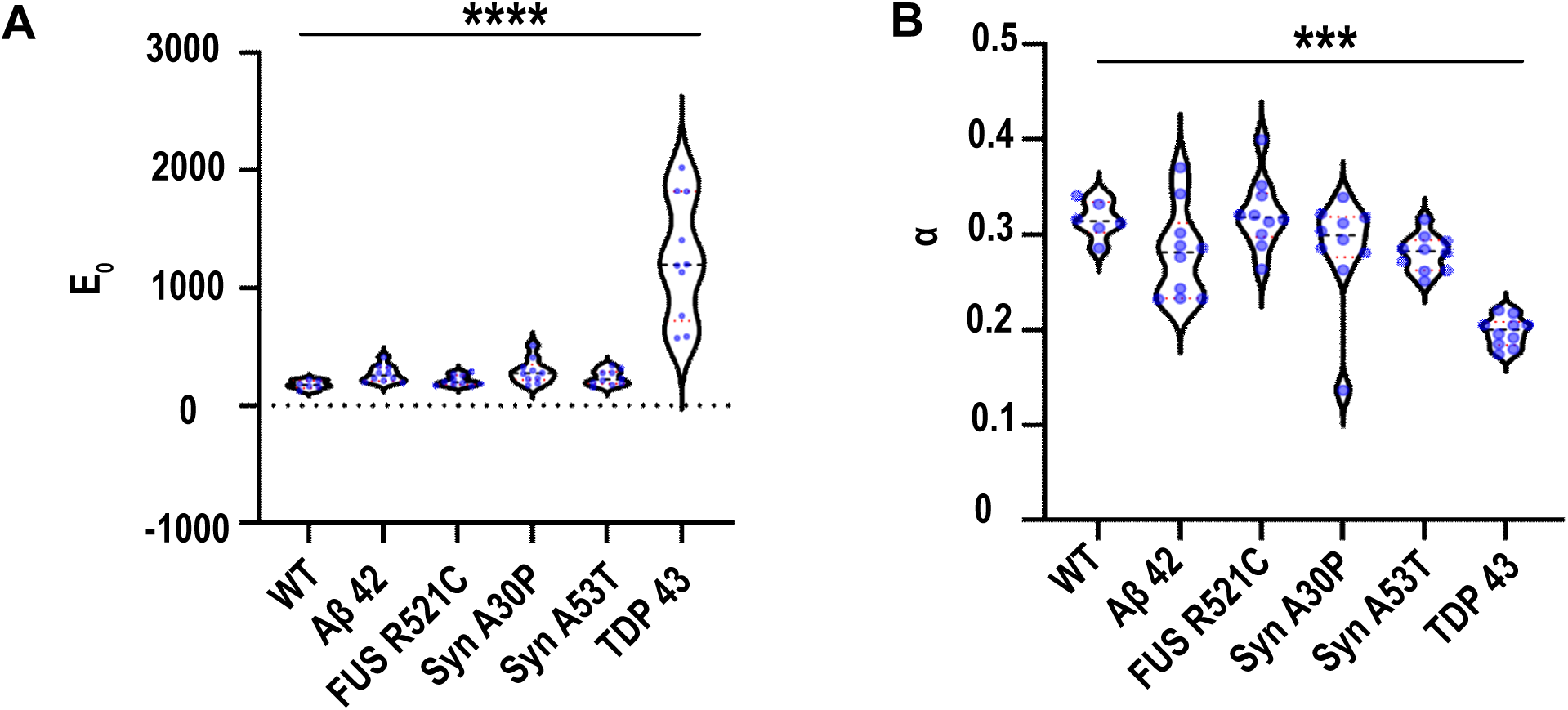
**A)** Box plot showing stiffness of all the cell types characterized by E_0._ WT (n=6), Aβ42 (n=10), FUS (n=10), HTT Q138 (n=10), αSynA30P (n=10), αSynA53T (n=10) and TDP43 (n=10) expressing cells. ****, P value < 0.0001. Statistical significance was determined using Kruskal-Wallis test followed by *post-hoc* Dunn’s multiple comparisons. B) Graph shows the α value for wild type cells and cells expressing various aggregate forming proteins. WT (n=6), Aβ42 (n=10), FUS (n=10), HTT Q138 (n=10), α-SynA30P (n=10), α-SynA53T (n=10) and TDP43 (n=10) expressing cells. ***, P value = 0.0001.

**Fig.9.**
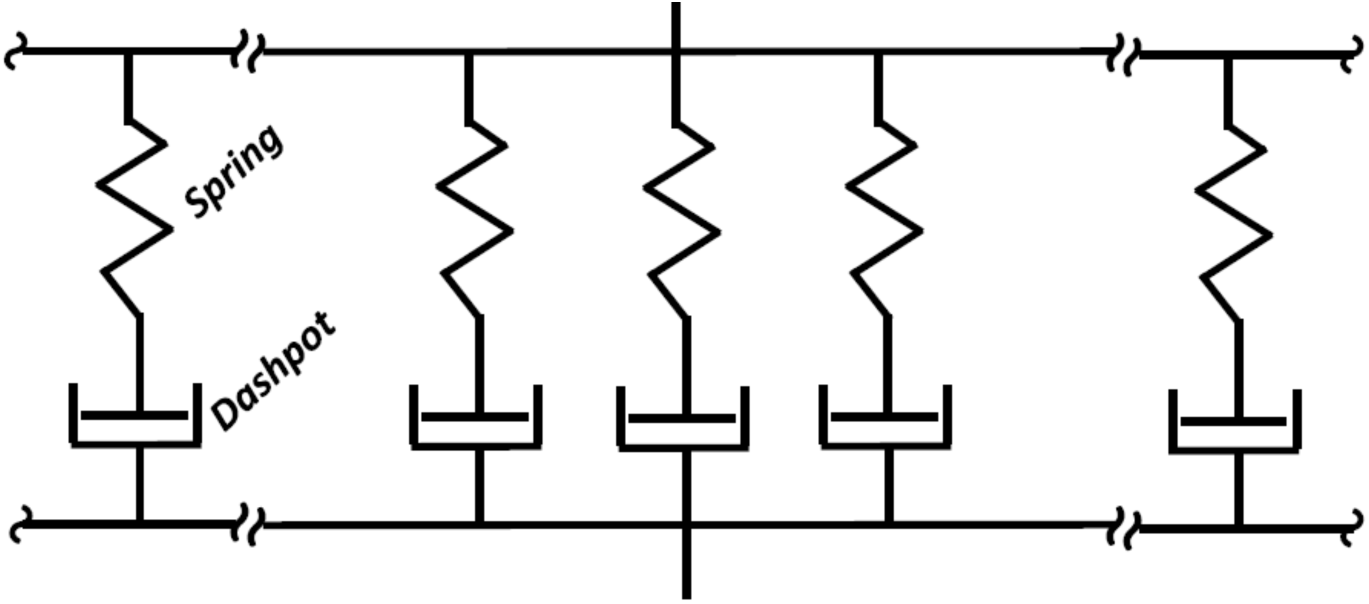
Schematic representation of PLR viscoelastic material. An infinite number of Maxwell elements-a series combination of a spring and a dashpot-are connected in parallel. This arrangement shows a continuous relaxation of the material.

In our study, we speculate that the altered physical properties of cells are directly linked to actin dynamics and hence the CME process. The PIV analysis of various cells and their phenotypes (WT, HTTQ138 aggregate, and rescue) suggest that actin dynamics in the presence of HTT138Q and TDP-43 were restricted compared to WT cells, leading to compromised CCSs movement (Fig. 1 and Fig. 6). The cells with HTT138Q and TDP-43 aggregates, were observed to have increased E_0_ and reduced α values compared to WT which suggests stiffening of cells accompanied by loss of fluidity. This is also observed to be the result of altered actin dynamics. Recovery of E_0_ and α values of cells upon overexpression of actin binding proteins (Hip1 and Arp2/3) reverts cells towards the WT, providing strong support to our hypothesis. More importantly, our AFM results show that rheological response of cells can be used as a biomarker for actin mediated altered cell behavior in the presence of pathogenic protein aggregates. Further, our results demonstrate that the toxicity caused by pathogenic protein aggregates may be due to a reorganization of other proteins, organelles and cellular pathways rendering them dysfunctional. Rescuing some of these functions may indeed work towards reducing neurotoxicity even in the continued presence of pathogenic aggregates.

## Discussion

Endocytosis has been shown to be affected in the context of neurodegeneration caused by pathogenic aggregates. However, it is as yet unclear as to how exactly such aggregates alter the process of endocytosis. Our work assesses CME by tracking clathrin light chain-positive structures in live cells in the presence of aggregating proteins responsible for neurodegeneration. The extent of observed change in CCS speed and directionality varied depending on the mutant protein present, suggesting that specific interactions may be responsible for the phenotype, rather than merely the physical presence of an aggregate in the cell. The most drastic changes resulting in a complete block in CCS movement were observed in the context of HTTQ138 and TDP-43. While the speed of CCS movement was also marginally altered in the presence of other mutant proteins, a complete block was only observed in the presence of HTTQ138 and TDP-43 (Fig. 1 and Fig. 6). Furthermore, these alterations in CCS movement also correlated with changes in actin dynamics as reflected by changes in filopodia length and lifetimes (Fig. 3 and Fig. 7). Inhibition of polymerization of the actin cytoskeleton either by chemicals (LatA/ CytoD) or by genetic means (profilin knockdown) also led to stalled CCSs movement, similar to what was observed in the presence of HTTQ138 or TDP-43 aggregates. Our results also revealed that Arp2/3 was sequestered by HTTQ138 aggregates (Fig. 3), and that overexpression of Arp2/3 rescued CCSs movement and filopodia formation even in presence of HTTQ138 aggregates (Fig. 4). Conversely, inhibiting the function of Arp2/3 either by CK666, (Supplementary Fig. 2) or by RNAi resulted in stalled CCS movement (Fig. 2), indicating that the branching function of Arp2/3 is essential for CME, and that the altered function of Arp2/3 by HTTQ138 aggregates may be one of several possible mechanisms by which CME is inhibited in Huntington’s disease. CCS movement in the presence of HTTQ138 aggregates could also be restored by overexpression of Hip1 or the chaperone, Mrj (Fig. 4).

Previous reports have demonstrated a tight correlation between the availability/distribution of Hip1R, Arp2/3 complex and endocytic rate and efficiency^62^. As the available number of Arp2/3 molecules increase, internalization depth increased in a shorter span of time. Further, while the distribution of Arp2/3 did not affect endocytic efficiency, the distribution of Hip1R regulated the endocytosis rate and extent, presumably by regulating the recruitment and distribution of Arp2/3 complexes on the surface of the growing endocytic pit. In vitro experiments have also revealed that the ratio of branching to capping protein regulates stiffness of the actin network. Network stiffness increased as either capping or branching protein concentration was increased^63^. The authors observed that this was in contrast to what was predicted, as increased capping should result in shorter filaments which should result in a softer network. However, this points to the fact that network stiffness is dependent on regulating a balance between branching and capping, and that an alteration in these may affect stiffness and endocytic efficiency.

FRAP of the clathrin light chain reveals that considerable exchange exists between the membrane associated and soluble pools of the protein even under endocytosis-blocking conditions^64, 65^. FRAP of the clathrin light chain in the presence of HTTQ138 and TDP-43 aggregates showed an incomplete recovery at CCSs (Supplementary Fig. 2,4). Furthermore alteration of actin cytoskeleton also shows delayed recovery of clathrin (Supplementary Fig. 2). These results indicate that availability/exchange of clathrin at the sites of CME is compromised in both these cases, in a manner hitherto unknown.

Our results suggest that the changes observed in CME in the context of HTTQ138 and TDP-43 may be dependent on the actin cytoskeleton, whereas this may not be the case in the context of other aggregates. Finally, we also determine that the mechanical response as measured by cellular stiffness is also significantly altered in the presence of HTTQ138 and TDP-43 (Fig. 5,8). Previous work from our lab has demonstrated that a loss of CME in embryonic stem cells results in an altered organization of the actin cytoskeleton and an increased cellular stiffness^20^. The mechanical response by measuring α, observed in the presence of HTTQ138 and TDP-43 aggregates also appears to be linked to the altered and less dynamic state of actin organization. The branched actin network is known to generate force required in many cellular processes including endocytosis, with alterations in this network directly limiting force transmission^66–68^. Our results show that regulating the cells’ physical properties, which is principally due to the actin network stiffness, by overexpressing Hip1 or Mrj, or by transient treatment with LatA could restore the stiffness of cells to WT levels even in presence of HTTQ138 aggregates. This suggests that a ‘Goldilocks’ state of the branched actin network is required for sufficient and efficient force generation during CME.

Why do various aggregating pathogenic proteins display such a variety of phenotypes with respect to CME and actin organisation? The presence of such altered and varied mechanisms have been reported even in the context of a single pathogenic protein. Overexpression of the monomeric versus the dimeric form of α-synuclein in the lamprey giant reticulospinal synapses demonstrated a block of CME at different stages indicative of their ability to interfere at different stages of CME, suggestive of a differential interaction with proteins^69^. TDP-43 knockdown has also been shown to alter the number and motility of Rab11-positive recycling endosomes^70^, while the loss of HTT has been shown to result in the altered activation status of cofilin, thereby resulting in altered actin dynamics^21^.

Recent work has highlighted a role for Nwk/FCHSD2, Dap160/Intersectin and WASP in regulating actin assembly at synapses^71^. Interestingly, Intersectin has been shown to associate with mutant forms of HTT and enhance its aggregation resulting in increased neuro-toxicity^72^. It remains unknown whether TDP-43 also exhibits similar interactions, thus resulting in similar phenotypes with respect to CCS dynamics and cellular stiffness. Additionally, it is also unknown whether other mutant proteins involved in aggregation and neurodegeneration interact with specific proteins involved in actin organization and endocytosis at the synapse, which in turn may affect their aggregation. These would form the basis for future studies and may also shed light on whether the endocytic phenotypes in these aggregate-containing cells are due to similar or different molecular mechanisms.

## Materials and Methods

### Drosophila strains and crosses

*Drosophila* strains were obtained from Bloomington Drosophila Stock Centre: Cg GAL4 (BL 27396), UAS-Clc-GFP (BL 7107), UAS-Lifeact-GFP (BL 56842), UAS-SNCA-A30P (Synuclein) (BL 8148), UAS-SNCA-A53T (BL 8147), TRiP.HMS00550 (BL 34523, Profilin RNAi), TRiP.HMS01846 (BL 38377, Hip1 RNAi), UAS-hHip1-HA (BL 66279), UAS-Arp3-GFP (BL 39722). ARP3-RNAi (VDRC# 35260) was obtained from Vienna Drosophila Resource Centre. UAS-Mrj-HA was generated in AM lab at NCCS. UAS-FUS-R521C and UAS-TDP-43 WT strains were a kind gift from Udai Bhan Pandey (University of Pittsburgh School of Medicine). Fly strain UAS - Amyloid β-42 - 2x / CyO was a kind gift from Diego E. Rincon Limas and Pedro Fernandez Funez (McKnight Brain Institute, University of Florida). UAS-HTTQ138-RFP was a gift from Troy Littleton (Massachusetts Institute of Technology).

Recombinant fly lines were generated by crossing CgGAL4 flies with Clathrin light chain (Clc) tagged with GFP (UAS-Clc-GFP) or with Lifeact tagged with GFP (UAS-Lifeact-GFP) flies. GFP positive larvae were selected and flies emerging from these larvae were balanced with CyO and maintained as recombinant lines for further use. These were crossed with aggregate protein-expressing flies, and hemocytes from third instar larvae were collected.

### Hemocyte isolation

Hemocytes used for all experiments were isolated from third instar larvae. Single larvae were dissected with the help of fine tweezers, in Schneider’s medium (S2 cell medium) and hemocytes were collected in a 35 mm glass bottom dish.

### mBSA internalization assay

Maleylation of Bovine serum Albumin (BSA, MP biomedicals) was performed as described earlier^73^. Fluorescent tagging of mBSA was performed using Alexa Fluor 594 Microscale Protein Labelling kit (Invitrogen, Molecular Probes) as per manufacturer’s protocol. For mBSA uptake, larvae were dissected and hemocytes were collected in serum free Schneider’s Drosophila medium. Cells were incubated for 10 – 15 min at room temperature to allow them to settle down. Cells were then incubated with mBSA at a concentration of 1 µg/ml for 35 min and washed several times with ice cold PBS. Fixation of hemocytes was done using 4% PFA for 20 min and cells were washed before imaging.

#### Inhibitor treatment

Hemocytes were treated with the following inhibitors at the mentioned concentrations: Latrunculin A (Sigma Aldrich cat. no. L5163) and Cytochalasin D (Sigma Aldrich, cat no. C8273), were used at a final concentration of 1µM. Latrunculin A was added at the time of imaging, while cells were treated for 1 hour with Cytochalasin D prior to imaging. CK666 (Sigma – Aldrich, cat. no. SML0006), was used at a final concentration of 50 µM, and cells were treated for 40 minutes prior to imaging. SMIFH2 (Sigma – Aldrich, cat. No. 344092), was used at a final concentration of 50 µM and cells were incubated for 15 minutes at room temperature before time lapse imaging.

### Imaging and image analysis

Time lapse imaging of hemocytes were performed along a single X-Y plane close to the glass coverslip to look at clathrin light chain and actin dynamics. Imaging was carried out for five minutes at five second intervals using a Nikon Ti Eclipse microscope. All images were acquired at 2.6X zoom using 100X plan Apo objective with 1.49 NA, using manufacturer’s software. To perform FRAP, hemocytes were isolated in S2 medium in glass-bottomed confocal dishes. ROI was drawn on a single non-motile CCS at the periphery of cells and photo-bleached using 10% laser power of the 488-nm laser with 100X, 1.49 NA oil immersion objective and pinhole of 1.2 AU. Pre-bleach time lapse images were acquired for 15 frames at 5.66 sec interval followed by bleaching for one frame for 0.971 sec duration. To look at the recovery of Clc-GFP, CCSs were then imaged for 50 frames at 5.66 sec interval. FRAP analysis was done using the online easyFRAP tool^74^ and mean intensity from five independent experiments was plotted.

Kymographs were generated using Image J software. CCSs were tracked using the Low Light Tracking Tool plugin in Fiji/ImageJ^75^. The resulting tracks were analysed and quantified using custom functions written in MATLAB, Mathworks. The mean instantaneous velocities of CCSs were quantified by calculating the distance moved by an individual CCS for each 5s interval of the time-lapse images, and averaging these instantaneous velocities for that CCS. All plots were generated in MATLAB. For tests of significance, data were first checked for normality using the chi2gof function and MATLAB.

Length and number of filopodia were quantified using Filoquant software, keeping the threshold constant across all the cells. Lifetimes of filopodia were calculated by manually calculating the duration that a filopodia persisted in a cell.

PIV analysis for actin flow was performed as previously described^76^ using the following PIV code : HTTps://github.com/stemarcotti/PIV. Source size was set to 0.5µm, while search size was set to 1.0µm. All other parameters were kept the same as in the given code. PIV analysis was performed on atleast 5 cells in each case.

### Quantification of 2D flow-fields from the time-lapse movies

To understand the movement of CCSs, we computed the 2D velocity flow-field 𝑣(𝑥, 𝑦) by performing a Particle Image Velocimetry (PIV)^77^ analysis on the time-lapse movies obtained from the experiments. Prior to the PIV analysis, the movies were corrected for drift errors using StackReg, an ImageJ plugin, after which an intensity correction (see Supplementary material for details) was applied to account for the bright background clathrin intensity which could significantly modify the nature of the flow-field.

### PIV analysis of the corrected time-lapse movies

The PIV analysis was performed using the PIV lab toolbox via Matlab (HTTps://pivlab.blogspot.com/). Before running the PIV analysis, the region-of-interest was chosen around the cell boundary and a common mask was applied to each frame to ensure that there was no spurious intensity from regions outside the cell boundary. To calculate the cross-correlation during the PIV analysis, we chose the Fast Fourier Transform (FFT) window deformation, with three passes starting from 512 pixels and with the final interrogation area of 64 pixels. After performing the PIV analysis over each movie, we performed a post-processing over each time-frame. First we applied a standard deviation filter with threshold value of 4 followed by a local median filter with threshold value 4. This ensured that we excluded velocity vectors with large variabilities. We also apply the interpolation filter which returns interpolated vector values for the missing data. The total interpolated vectors per frame ∼10%. Finally, we also applied a one-step spatial smoothing of the velocity flow-field for each frame. Note that we have checked our results by performing the PIV analysis (i) without applying the interpolation filter at each frame, and, (ii) with a finer binning where the final interrogation area taken to be 32 pixels.. For both these cases we obtain a similar radial flow profile 𝑉_*r*_(𝑟) (discussed in the following sections). We note from the radial speed profiles that for the Wild Type cells, the maximum radial speed, which is attained near the cell membrane is 𝑉_*r*_ 𝑟 → 𝑅 ≈ 0.005 𝜇𝑚/𝑠𝑒𝑐. This value is lower than the speed values estimated from the particle tracking experiments (Fig. 6C). This is not surprising as the PIV analysis gives an estimate of the hydrodynamic flow-fields by computing the average displacement of a large number of particles. On the hand other individual particle tracking is more precise and was done over selective trajectories taken close to the membrane and also across a large number of cells. Thus it is expected that PIV analysis will underestimate the speed magnitudes compared to the original value and the degree of which depends on the presence of spurious noise in the movies. However on the other hand, PIV analysis efficiently captures the large-scale continuum flow-field and may reveal more about the underlying physical principles that lead to such collective flows.

### Time-averaged steady-state flow profile

To further quantify the flow profiles from the PIV analyzed movies, we made a steady-state assumption, since the overall flow pattern did not change over the duration of movies.

Under the steady-state assumption, we obtain the time-averaged spatial flow-field profile by averaging over the entire time duration 𝑇 = 𝑁𝛥𝑡 of the movie:

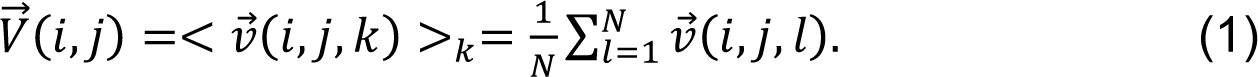

Here 𝑣(𝑖, 𝑗, 𝑘), is the flow-vector, where each (𝑖, 𝑗) value indicate the pixel coordinates, and the index 𝑘 indicates the time-frame. Here 𝑁 is the number of time-frames and 𝛥𝑡 is the inverse frame-rate. Finally using the pixel to micron conversion information (Supp. Table 1), we obtain the average spatial flow-field 𝑉(𝑥, 𝑦) where (𝑥, 𝑦) = (𝑖𝛥𝑥, 𝑗𝛥𝑦). In supplementary figure 1b, we show the time-averaged flow-field obtained from PIV analysis plotted on top of the corrected intensity field for the wild type cell.

### Quantification of PIV data

From the time-averaged spatial flow-field profile for the Wild type (Supplementary fig. 1b), it is evident that the CCSs flow radially inwards from the cell membrane to the cell center while in the mutants such as HTTQ138 and cells treated with LAT-A, this movement is severely affected. To quantify the nature of this radial movement and to compare between the different experimental cases, we obtained the radial speed profile from the time-average spatial flow field 𝑥, 𝑦 . As a first step, we identified the cell centre (𝑥_*c*_, 𝑦_*c*_). The flow-field was then translated to new coordinates 𝑥 → (𝑥 − 𝑥_*c*_), 𝑦 → (𝑦 − 𝑦_*c*_). We then performed a transformation to polar coordinates (𝑟, 𝜃), where 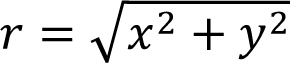, and 𝜃 =𝑡𝑎𝑛^−1^ (𝑦/𝑥) and computed the radial component of the speed, which is given as 𝑉_*r*_(𝑟, 𝜃) = 𝑉(𝑥, 𝑦) ⋅ 𝑟 = (𝑉_*r*_ 𝑐𝑜𝑠 𝜃 + 𝑉_*r*_ 𝑠𝑖𝑛 𝜃). Note that one can also similarly compute the polar component 𝑉_*r*_(𝑟, 𝜃) which is negligible for the Wild type, and therefore we do not quantify it. Next we obtained the radial flow profile purely as a function of the radial distance 𝑟, namely 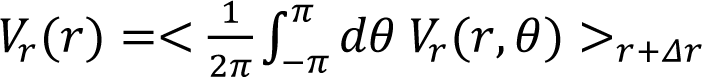. Note that the 𝑉_*r*_(𝑟) value was obtained by averaging over all possible 𝜃 values over an annulus between [𝑟, 𝑟 + 𝛥𝑟]. We choose 𝛥𝑟 = 60 pixels such that 0.5𝜇𝑚 ≤ 𝛥𝑟 ≤ 1.2𝜇𝑚 (see Supp. Table 1). Note that to obtain 𝑉_*r*_(𝑟), we assumed that the flow-field has a polar symmetry. While the cells are not exactly circular and a more rigorous quantification in that case would be to estimate the exact cell boundary contour and perform the polar-angle averaging over concentric contours. However, the nature of the flow-field will remain unaffected and we therefore proceeded with this simplified definition.

Finally we quantified the directionality of the flow-field by computing the distribution of the velocity direction relative to the polar direction. To obtain this distribution, we first constructed the variable 𝜃_*dir*_ = (𝜃_*v*_ − 𝜃), where 𝜃_*v*_ =𝑡𝑎𝑛^−1^ (𝑉_*r*_/𝑉_*r*_). For a pure radial flow, we will have either 𝜃_*dir*_ = 0 or 𝜃_*dir*_ = 𝜋, depending on whether the flow is radially inward or outward. For purely circular motion, 𝜃_*dir*_ = ±𝜋/2, depending on whether it is clockwise or counter-clockwise. Once 𝜃_*dir*_ is obtained for each pixel (𝑟, 𝜃), we compute the corresponding normalized histogram 𝑃(𝜃_*dir*_).

#### S2 cell culture and transfection

Approximately 1× 10^6^ cells were plated in 12 well plates in S2 cell medium (Schneider’s medium). Cells were transfected with Huntingtin mRFPQ15, Huntingtin mRFPQ138 and Actin-GFP plasmid along with PAC-GAL4 vector at a concentration of 200ng each, using PEI (Polyethyleneimine). S2 cells were grown at 25^ᵒ^C in a humidified incubator in 5% CO_2._

#### FRAP and analysis

To perform FRAP, hemocytes were isolated in S2 medium in glass bottom plates. ROI was drawn on single non-motile CCS at the periphery of cells and photo-bleached using 10% laser power of 488-nm laser with 100X, 1.49 NA oil immersion objective and pinhole of 1.2 airy unit. Pre-bleached time lapse images were acquired for 15 frames at 5.66 sec interval followed by bleaching for one frame for 0.971 sec duration. To look at the recovery of Clc-GFP, CCSs were imaged for 50 frames at 5.66 sec interval. FRAP analysis was done using the online easyFRAP tool^74^ and mean intensity from five independent experiments was plotted using Graphpad Prism.

### Atomic force microscopy and nanoindentation experiments

The nanoindentation experiments on single cells were performed using JPK Nanowizard II AFM from Berlin, Germany. The tipless cantilevers from MikroMasch, Bulgaria (model no.- HQ:CSC38/tipless/Ce-Au) were used. The tipless commercial cantilever was customized by attaching a 5 µm (diameter) glass-bead using lift-off method as described previously^20^. Briefly, the resin and hardener of a two-component epoxy (Araldite Klear, India) were mixed in 1:1 ratio and stirred until the mixture colour turns gray. A few tiny glue-droplets were decorated on a clean glass slide. The cantilever was approached manually on one of the droplets carefully such that the cantilever’s free end should just touch the droplet’s topmost layer and pick a tiny amount of glue, while avoiding dipping the cantilever. The cantilever was retracted by ∼ 20 µm. It was brought near the area where the glass-beads were sparsely coated on the same glass slide. The cantilever was brought into contact on a clean area several times at different locations on the glass slide to reduce the excess amount of glue. The cantilever was brought onto a clean glass bead and aligned in such a way that the bead would attach at the desired position. Once aligned, it is approached using the autoapproach mechanism with the setpoint of 0.15-0.2 Volts. Once the autoapproach was done, the setpoint was gradually increased to 1.0 Volt. The glue is allowed to cure for 20-30 minutes. The cantilever was retracted and it was confirmed that the bead remained attached. The position of bead attachment was determined by Scanning Electron Microscope (SEM) imaging. We only used the cantilevers with beads attached in the middle along the cantilever width.

The photodetector sensitivity and cantilever force constant were determined before every experiment. The detection sensitivity (in nm/V) was determined by obtaining a slope of approach curve at the cantilever-glass slide deep contact region. The force constant was determined using thermal tuning method^78^ available in the software. The cantilevers with a typical force constant of 0.2 N/m were used for experiments.

#### Nanoindentation Experiment

Hemocytes were isolated from third instar larvae and plated onto Concanavalin-coated 22mm circular glass coverslips, which were freshly cleaned and wiped with methanol to avoid unwanted dirt on the surface before experiments. Cells were incubated for 30 minutes at room temperature to allow proper adherence to the coverslip. The approach-retract experiments were performed on a single cell with ∼ 2 µm/s speed on a 1 x 1 µm^2^ area using a 7 x 7 grid with a sampling rate of 2 kHz. We allowed a rest period of 2 seconds between two successive force curves for the cell to recover its original shape. The experimentally recorded parameters (raw data) were cantilever deflection (*d_cant_*, in Volts) and the base piezo displacement (*d_pz_*, in Volts). The piezo displacement corresponds to movement of the cantilever base. *d_cant_* and *d_pz_* were converted to the unit of meters by multiplying them with photodetector sensitivity and base-piezo sensitivity, respectively. The force (*F*) on the cantilever/sample was calculated by multiplication of *d_cant_* with the cantilever force constant. The cell deformation was determined by subtracting cantilever deflection from the base-piezo displacement.

#### Data Analysis

The force curves were fitted with Ting’s model and the cell’s viscoelastic properties were extracted. We followed the procedure developed by Y. Effremove et al^61^. According to Ting’s model, the relation between force and deformation of a linear viscoelastic material is as follows:

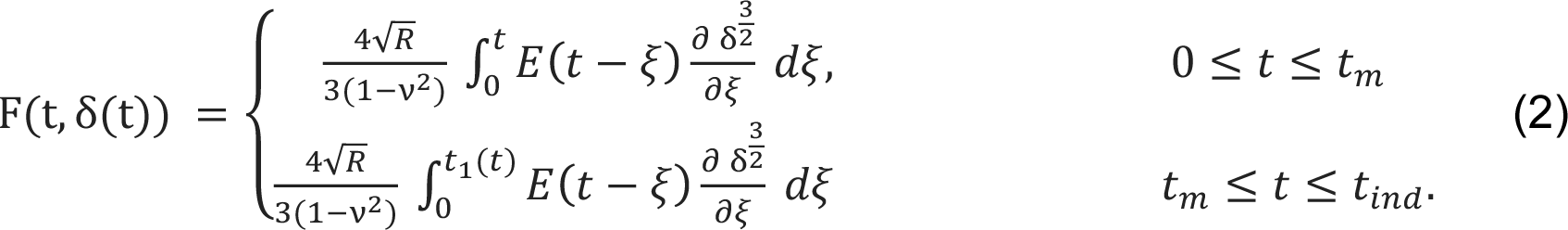

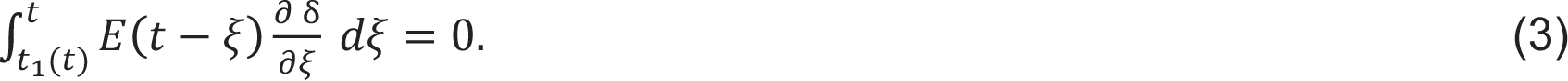

Where 𝑅 and ν are the radius of the indenter and Poisson’s ratio of the sample respectively. ξ is the dummy time variable used for the integration. 𝐸 𝑡 is the relaxation modulus and can be used from any linear viscoelastic constitutive equations that better describe the sample’s behavior. We found that the power-law rheology (PLR) best describes the cell’s viscoelastic behavior as also reported by Efremov et al^61^ for the fibroblast cells. In the PLR model, the viscoelasticity is modeled with an infinite number of Maxwell components connected in parallel which show a continuous relaxation and power law decay. The relaxation modulus for the modified power-law rheology (mPLR) model is as follows:

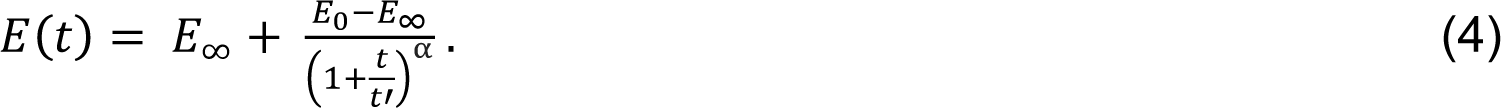

Where 𝐸_0_, 𝐸_∞_, and *α* are the instantaneous, long-term Young’s modulus and power law exponent respectively. 𝑡′ is a small time offset and it is set to be sampling time (inverse of the sampling rate). We found 𝐸_∞_∼ 0 for the cells (data not shown), so we employed the simple power-law rheology (or PLR) model. The relaxation modulus for PLR is following-

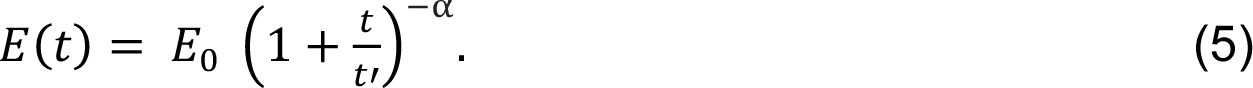

The use of the above-simplified form also reduces the program run time during the analysis. The above-simplified form of the relaxation modulus helps to reduce the program run time during the analysis.

#### Author contributions

SBS, DS and AM conceptualized the project. SBS carried out all experiments described in Fig 1-7, Supp. Fig.1,2,4. AS and AN conducted PIV analysis on data described in Fig. 1,2,4,Supp Fig.1. VA analysed data in Fig 6,7. DS analysed data in Fig. 3E. SR and SP carried out AFM experiments described in Fig. 5,8, Supp.Fig. 3,5. AM conducted experiments described in Fig. 4G. All authors helped write the first draft of the manuscript and reviewed subsequent versions.

## Supporting information

SuppMovie 2B

## Acknowledgements

This work was supported by funding to DS and AM from an NCCS Intramural Collaborative grant NCCS/DIR/2018/24. SP acknowledges funding support from Department of Science and Technology, Govt. of India (CRG/2022/001891). AM acknowledges funding from Wellcome Trust-DBT India Alliance (IA/I/13/2/501030), and Department of Biotechnology, Govt of India (BT/PR25893/GET/119/174/2017). VA is supported by funding from EMBL Australia. We thank Stefania Marcotti for sharing the PIV code. We thank UB Pandey, DE Rincon Limas, PF Funez and T. Littleton for sharing fly lines. We thank Abhilipsa Das and Shreejita Chatterjee for help with experiment 4G.

## Conflict of Interest

The authors declare no conflict of interest.

## Notes

### Competing Interest Statement

The authors have declared no competing interest.

## References

1. DiFiglia, M. et al. Aggregation of huntingtin in neuronal intranuclear inclusions and dystrophic neurites in brain. Science 277, 1990–1993 (1997).

2. Chartier-Harlin, M.-C. et al. Early-onset Alzheimer’s disease caused by mutations at codon 717 of the β-amyloid precursor protein gene. Nature 353, 844–846 (1991).

3. Bruijn, L. et al. ALS-linked SOD1 mutant G85R mediates damage to astrocytes and promotes rapidly progressive disease with SOD1-containing inclusions. Neuron 18, 327–338 (1997).

4. Davies, S. W. et al. Formation of neuronal intranuclear inclusions underlies the neurological dysfunction in mice transgenic for the HD mutation. Cell 90, 537–548 (1997).

5. Wilson, D. M. et al. Hallmarks of neurodegenerative diseases. Cell 186, 693–714 (2023).

6. Reiner, A. et al. Differential loss of striatal projection neurons in Huntington disease. Proc. Natl. Acad. Sci. 85, 5733–5737 (1988).

7. Lin, C.-H. et al. Neurological abnormalities in a knock-in mouse model of Huntington’s disease. Hum. Mol. Genet. 10, 137–144 (2001).

8. Strong, T. V. et al. Widespread expression of the human and rat Huntington’s disease gene in brain and nonneural tissues. Nat. Genet. 5, 259–265 (1993).

9. MacDonald, M. et al. The Huntington’s Disease Collaborative Research Group. A novel gene containing a trinucleotide repeat that is expanded and unstable on Huntington’s disease chromosomes. Cell 72, 971–983 (1993).

10. Lunkes, A. et al. Properties of polyglutamine expansion in vitro and in a cellular model for Huntington’s disease. Philos. Trans. R. Soc. Lond. B. Biol. Sci. 354, 1013–1019 (1999).

11. Krobitsch, S. & Lindquist, S. Aggregation of huntingtin in yeast varies with the length of the polyglutamine expansion and the expression of chaperone proteins. Proc. Natl. Acad. Sci. 97, 1589–1594 (2000).

12. Babcock, D. T. & Ganetzky, B. Transcellular spreading of huntingtin aggregates in the Drosophila brain. Proc. Natl. Acad. Sci. 112, E5427–E5433 (2015).

13. Gauthier, L. R. et al. Huntingtin controls neurotrophic support and survival of neurons by enhancing BDNF vesicular transport along microtubules. Cell 118, 127–138 (2004).

14. Yu, A., et al. Protein aggregation can inhibit clathrin-mediated endocytosis by chaperone competition. Proc. Natl. Acad. Sci. 111, E1481–E1490 (2014).

15. Royle, S. J. & Lagnado, L. Clathrin-­-mediated endocytosis at the synaptic terminal: bridging the gap between physiology and molecules. Traffic 11, 1489– 1497 (2010).

16. Conner, S. D. & Schmid, S. L. Regulated portals of entry into the cell. Nature 422, 37–44 (2003).

17. Kaltenbach, L. S. et al. Huntingtin interacting proteins are genetic modifiers of neurodegeneration. PLoS Genet. 3, e82 (2007).

18. Meriin, A. B. et al. Aggregation of expanded polyglutamine domain in yeast leads to defects in endocytosis. Mol. Cell. Biol. 23, 7554–7565 (2003).

19. Merrifield, C. J., Feldman, M. E., Wan, L. & Almers, W. Imaging actin and dynamin recruitment during invagination of single clathrin-coated pits. Nat. Cell Biol. 4, 691–698 (2002).

20. Mote, R. D. et al. Pluripotency of embryonic stem cells lacking clathrin-mediated endocytosis cannot be rescued by restoring cellular stiffness. J. Biol. Chem. 295, 16888–16896 (2020).

21. Wennagel, D., Braz, B. Y., Capizzi, M., Barnat, M. & Humbert, S. Huntingtin coordinates dendritic spine morphology and function through cofilin-mediated control of the actin cytoskeleton. Cell Rep. 40, (2022).

22. Weiss, K. R., Kimura, Y., Lee, W.-C. M. & Littleton, J. T. Huntingtin aggregation kinetics and their pathological role in a Drosophila Huntington’s disease model. Genetics 190, 581–600 (2012).

23. Doornaert, B. et al. Time course of actin cytoskeleton stiffness and matrix adhesion molecules in human bronchial epithelial cell cultures. Exp. Cell Res. 287, 199–208 (2003).

24. Bhadriraju, K. & Hansen, L. K. Extracellular matrix-and cytoskeleton-dependent changes in cell shape and stiffness. Exp. Cell Res. 278, 92–100 (2002).

25. Tavares, S. et al. Actin stress fiber organization promotes cell stiffening and proliferation of pre-invasive breast cancer cells. Nat. Commun. 8, 15237 (2017).

26. Mettlen, M. & Danuser, G. Imaging and modeling the dynamics of clathrin-mediated endocytosis. Cold Spring Harb. Perspect. Biol. 6, a017038 (2014).

27. Kochubey, O., Majumdar, A. & Klingauf, J. Imaging clathrin dynamics in Drosophila melanogaster hemocytes reveals a role for actin in vesicle fission. Traffic 7, 1614–1627 (2006).

28. Yarar, D., Waterman-Storer, C. M. & Schmid, S. L. A dynamic actin cytoskeleton functions at multiple stages of clathrin-mediated endocytosis. Mol. Biol. Cell 16, 964–975 (2005).

29. Ferguson, S. et al. Coordinated actions of actin and BAR proteins upstream of dynamin at endocytic clathrin-coated pits. Dev. Cell 17, 811–822 (2009).

30. Korobova, F. & Svitkina, T. Arp2/3 complex is important for filopodia formation, growth cone motility, and neuritogenesis in neuronal cells. Mol. Biol. Cell 19, 1561–1574 (2008).

31. Rouiller, I. et al. The structural basis of actin filament branching by the Arp2/3 complex. J. Cell Biol. 180, 887–895 (2008).

32. Volkmann, N. et al. Structure of Arp2/3 complex in its activated state and in actin filament branch junctions. science 293, 2456–2459 (2001).

33. Mullins, R. D., Heuser, J. A. & Pollard, T. D. The interaction of Arp2/3 complex with actin: nucleation, high affinity pointed end capping, and formation of branching networks of filaments. Proc. Natl. Acad. Sci. 95, 6181–6186 (1998).

34. Theriot, J. A. & Mitchison, T. J. The three faces of profilin. Cell 75, 835–838 (1993).

35. Suetsugu, S., Miki, H. & Takenawa, T. The essential role of profilin in the assembly of actin for microspike formation. EMBO J. 17, 6516–6526 (1998).

36. Subtil, A. & Dautry-Varsat, A. Microtubule depolymerization inhibits clathrin coated-pit internalization in non-adherent cell lines while interleukin 2 endocytosis is not affected. J. Cell Sci. 110, 2441–2447 (1997).

37. Rappoport, J. Z., Taha, B. W. & Simon, S. M. Movement of plasma-­-membrane-­-associated clathrin spots along the microtubule cytoskeleton. Traffic 4, 460–467 (2003).

38. Popova, J. S. & Rasenick, M. M. Clathrin-mediated Endocytosis of m3 Muscarinic Receptors: ROLES FOR Gβγ AND TUBULIN. J. Biol. Chem. 279, 30410–30418 (2004).

39. Caviston, J. P. & Holzbaur, E. L. Huntingtin as an essential integrator of intracellular vesicular trafficking. Trends Cell Biol. 19, 147–155 (2009).

40. Buss, F., Luzio, J. P. & Kendrick-Jones, J. Myosin VI, a new force in clathrin mediated endocytosis. FEBS Lett. 508, 295–299 (2001).

41. Soldati, T. & Schliwa, M. Powering membrane traffic in endocytosis and recycling. Nat. Rev. Mol. Cell Biol. 7, 897–908 (2006).

42. Buss, F., Arden, S. D., Lindsay, M., Luzio, J. P. & Kendrick-Jones, J. Myosin VI isoform localized to clathrin-coated vesicles with a role in clathrin-mediated endocytosis. EMBO J. 20, 3676–3684 (2001).

43. Wagner, W. et al. Myosin VI drives clathrin-mediated AMPA receptor endocytosis to facilitate cerebellar long-term depression. Cell Rep. 28, 11–20 (2019).

44. Biancospino, M. et al. Clathrin light chain A drives selective myosin VI recruitment to clathrin-coated pits under membrane tension. Nat. Commun. 10, 4974 (2019).

45. Metzler, M. et al. HIP1 functions in clathrin-mediated endocytosis through binding to clathrin and adaptor protein 2. J. Biol. Chem. 276, 39271–39276 (2001).

46. Engqvist-Goldstein, Å. E. et al. The actin-binding protein Hip1R associates with clathrin during early stages of endocytosis and promotes clathrin assembly in vitro. J. Cell Biol. 154, 1209–1224 (2001).

47. Kalchman, M. A. et al. HIP1, a human homologue of S. cerevisiae Sla2p, interacts with membrane-associated huntingtin in the brain. Nat. Genet. 16, 44–53 (1997).

48. Senetar, M. A., Foster, S. J. & McCann, R. O. Intrasteric inhibition mediates the interaction of the I/LWEQ module proteins Talin1, Talin2, Hip1, and Hip12 with actin. Biochemistry 43, 15418–15428 (2004).

49. Wilbur, J. D. et al. Actin binding by Hip1 (huntingtin-interacting protein 1) and Hip1R (Hip1-related protein) is regulated by clathrin light chain. J. Biol. Chem. 283, 32870–32879 (2008).

50. Labbadia, J. et al. Suppression of protein aggregation by chaperone modification of high molecular weight complexes. Brain 135, 1180–1196 (2012).

51. Warrick, J. M. et al. Suppression of polyglutamine-mediated neurodegeneration in Drosophila by the molecular chaperone HSP70. Nat. Genet. 23, 425–428 (1999).

52. Hageman, J. et al. A DNAJB chaperone subfamily with HDAC-dependent activities suppresses toxic protein aggregation. Mol. Cell 37, 355–369 (2010).

53. Kakkar, V. et al. The S/T-rich motif in the DNAJB6 chaperone delays polyglutamine aggregation and the onset of disease in a mouse model. Mol. Cell 62, 272–283 (2016).

54. Chuang, J.-Z. et al. Characterization of a brain-enriched chaperone, MRJ, that inhibits Huntingtin aggregation and toxicity independently. J. Biol. Chem. 277, 19831–19838 (2002).

55. Piette, B. L. et al. Comprehensive interactome profiling of the human Hsp70 network highlights functional differentiation of J domains. Mol. Cell 81, 2549–2565 (2021).

56. Desai, M. et al. Mrj an Hsp40 family chaperone regulates oligomerization of Orb2 and long-term memory. bioRxiv 2022–10 (2022).

57. Ruggeri, F. S., Šneideris, T., Vendruscolo, M. & Knowles, T. P. Atomic force microscopy for single molecule characterisation of protein aggregation. Arch. Biochem. Biophys. 664, 134–148 (2019).

58. Sollich, P. Rheological constitutive equation for a model of soft glassy materials. Phys. Rev. E 58, 738 (1998).

59. Kollmannsberger, P. & Fabry, B. Linear and nonlinear rheology of living cells. Annu. Rev. Mater. Res. 41, 75–97 (2011).

60. Kollmannsberger, P., Mierke, C. T. & Fabry, B. Nonlinear viscoelasticity of adherent cells is controlled by cytoskeletal tension. Soft Matter 7, 3127–3132 (2011).

61. Efremov, Y. M., Wang, W.-H., Hardy, S. D., Geahlen, R. L. & Raman, A. Measuring nanoscale viscoelastic parameters of cells directly from AFM force-displacement curves. Sci. Rep. 7, 1541 (2017).

62. Akamatsu, M. et al. Principles of self-organization and load adaptation by the actin cytoskeleton during clathrin-mediated endocytosis. Elife 9, e49840 (2020).

63. Pujol, T., du Roure, O., Fermigier, M. & Heuvingh, J. Impact of branching on the elasticity of actin networks. Proc. Natl. Acad. Sci. 109, 10364–10369 (2012).

64. Wu, X. et al. Clathrin exchange during clathrin-mediated endocytosis. J. Cell Biol. 155, 291–300 (2001).

65. Wu, X. et al. Adaptor and clathrin exchange at the plasma membrane and trans-Golgi network. Mol. Biol. Cell 14, 516–528 (2003).

66. Goode, B. L., Eskin, J. A. & Wendland, B. Actin and endocytosis in budding yeast. Genetics 199, 315–358 (2015).

67. Lacy, M. M., Ma, R., Ravindra, N. G. & Berro, J. Molecular mechanisms of force production in clathrin-­-mediated endocytosis. FEBS Lett. 592, 3586–3605 (2018).

68. Planade, J. et al. Mechanical stiffness of reconstituted actin patches correlates tightly with endocytosis efficiency. PLoS Biol. 17, e3000500 (2019).

69. Medeiros, A. T., Soll, L. G., Tessari, I., Bubacco, L. & Morgan, J. R. α-Synuclein dimers impair vesicle fission during clathrin-mediated synaptic vesicle recycling. Front. Cell. Neurosci. 11, 388 (2017).

70. Schwenk, B. M. et al. TDP-­-43 loss of function inhibits endosomal trafficking and alters trophic signaling in neurons. EMBO J. 35, 2350–2370 (2016).

71. Del Signore, S. J. et al. An autoinhibitory clamp of actin assembly constrains and directs synaptic endocytosis. Elife 10, e69597 (2021).

72. Scappini, E., Koh, T.-W., Martin, N. P. & O’Bryan, J. P. Intersectin enhances huntingtin aggregation and neurodegeneration through activation of c-Jun-NH2-terminal kinase. Hum. Mol. Genet. 16, 1862–1871 (2007).

73. Guha, A., Sriram, V., Krishnan, K. & Mayor, S. Shibire mutations reveal distinct dynamin-independent and-dependent endocytic pathways in primary cultures of Drosophila hemocytes. J. Cell Sci. 116, 3373–3386 (2003).

74. Koulouras, G. et al. EasyFRAP-web: a web-based tool for the analysis of fluorescence recovery after photobleaching data. Nucleic Acids Res. 46, W467– W472 (2018).

75. Krull, A. et al. A divide and conquer strategy for the maximum likelihood localization of low intensity objects. Opt. Express 22, 210–228 (2014).

76. Yolland, L. et al. Persistent and polarized global actin flow is essential for directionality during cell migration. Nat. Cell Biol. 21, 1370–1381 (2019).

77. Raffel, M., Willert, C. E. & Kompenhans, J. Particle Image Velocimetry [electronic resource]: A Practical Guide. (2007).

78. Heim, L.-O., Kappl, M. & Butt, H.-J. Tilt of atomic force microscope cantilevers: effect on spring constant and adhesion measurements. Langmuir 20, 2760–2764 (2004).

