## Supplementary information for "Pathogenic aggregates alter actin organization and cellular viscosity resulting in stalled clathrin mediated endocytosis"

**Supplementary Figure 1**: A) Micrograph showing sequestration of actin-GFP in HTT Q138 aggregates in Drosophila S2 cells. HTT Q15 or HTT Q138 is tagged with mRFP. HTT Q15 does not sequester actin and filopodia like projections are seen in HTT Q15 expressing cells. Scale bar 2µm. B) Time-averaged CCS flow-field obtained from the PIV analysis for Wild type cells. The flow-field vectors are plotted on top of the corrected clathrin intensity shown in gray-scale. The color of the vectors represent the vector magnitudes (see color-bar for the exact values in $\mu m/sec$.

**Supplementary Figure 2. A)** Kymograph showing stalled movement of CCSs upon treatment with CytoD, CK666, SMIFH2 and upon myosin VI knockdown. **B)** Graph showing fluorescence intensity of CLC GFP upon FRAP of individual CCSs. Y-axis depicts GFP intensity and X-axis represents time in seconds. Recovery of CLC GFP is seen at individual CCSs in WT and Luc VAL 10 (control for profilin KD). However, HTTQ138 expressing cells, Lat A treated cells and profilin knockdown cells only show partial recovery.

**Supplementary Figure 3. A)** A representative force curve obtained on wild type cells. The points (a), (b), (c), (d), and (e) represent the conditions at various phases of cell deformations while the bead indents on the cell, with the bead leaving the cell surface at point (e), compared to the point at which it makes contact while approaching (a).

**Supplementary Figure 4. A)** Graph showing fluorescence intensity of CLC GFP upon FRAP of individual CCSs. Y-axis depicts GFP intensity and X-axis represents time in seconds. Only partial recovery of CLC GFP is seen in presence of HTT Q138 and TDP-43 whereas recovery of CLC GFP in other aggregate-containing cells is comparable to WT cells.

**Supplementary Figure 5. A)** Representative force curves of each cell type fitted to Ting's model with Power Law Rheology.

**Supplementary movie 1A:** Video showing time lapse imaging of a HTTQ15 hemocyte expressing clathrin light chain tagged with GFP over 5 minutes at five second intervals.

**Supplementary movie 1B:** Video showing time lapse imaging of an HTTQ138 hemocyte expressing clathrin light chain tagged with GFP over 5 minutes at five second intervals.

**Supplementary movie 2A:** Video showing time lapse imaging of a WT hemocyte expressing Lifeact tagged with GFP over 5 minutes at five second intervals.

**Supplementary movie 2B:** Video showing time lapse imaging of a hemocyte expressing Lifeact tagged with GFP in presence of HTTQ138 over 5 minutes at five second intervals

**Supplementary movie 2C:** Video showing time lapse imaging of a WT hemocyte expressing Tubulin tagged with GFP over 5 minutes at five second intervals.

**Supplementary movie 2D:** Video showing time lapse imaging of a hemocyte containing HTTQ138 aggregates expressing Tubulin tagged with GFP over 5 minutes at five second intervals.

**Supplementary movie 3A:** Video showing time lapse imaging of a hemocyte expressing Clathrin light chain tagged with GFP over 5 minutes at five second intervals upon myosin VI knockdown condition.

**Supplementary movie 3B:** Video showing time lapse imaging of a hemocyte expressing Lifeact tagged with GFP over 5 minutes at five second intervals upon myosin VI knockdown condition.

**Supplementary movie 4A:** Video showing time lapse imaging of a hemocyte expressing Clathrin light chain tagged with GFP over 5 minutes at five second intervals under HTT Q138 + Arp2/3 condition.

**Supplementary movie 4B:** Video showing time lapse imaging of hemocyte expressing Lifeact tagged with GFP over 5 minutes at five second intervals under HTT Q138 + Arp2/3 condition.

**Supplementary movie 4C:** Video showing time lapse imaging of a hemocyte expressing Clathrin light chain tagged with GFP over 5 minutes at five second intervals under HTT Q138 + Hip1 condition.

**Supplementary movie 4D:** Video showing time lapse imaging of a hemocyte expressing Clathrin light chain tagged with GFP over 5 minutes at five second intervals under HTT Q138 + Mrj condition.

**Supplementary movie 5A.** Video showing time lapse imaging of a hemocyte containing Aβ42 aggregates expressing Clathrin light chain tagged with GFP over 5 minutes at five second intervals.

**Supplementary movie 5B.** Video showing time lapse imaging of a hemocyte containing FUS R521C aggregates expressing Clathrin light chain tagged with GFP over 5 minutes at five second intervals.

**Supplementary movie 5C.** Video showing time lapse imaging of a hemocyte containing α-SynA30P aggregates expressing Clathrin light chain tagged with GFP over 5 minutes at five second intervals.

**Supplementary movie 5D:** Video showing time lapse imaging of a hemocyte containing α-SynA53T aggregates expressing Clathrin light chain tagged with GFP over 5 minutes at five second intervals.

**Supplementary movie 5E:** Video showing time lapse imaging of a hemocyte containing TDP-43 aggregates expressing Clathrin light chain tagged with GFP over 5 minutes at five second intervals.

**Supplementary movie 6A:** Video showing time lapse imaging of a hemocyte containing Aβ42 aggregates expressing Lifeact tagged with GFP over 5 minutes at five second intervals.

**Supplementary movie 6B:** Video showing time lapse imaging of a hemocyte containing FUS R521C aggregates expressing Lifeact tagged with GFP over 5 minutes at five second intervals.

**Supplementary movie 6C:** Video showing time lapse imaging of a hemocyte containing α-SynA30P aggregates expressing Lifeact tagged with GFP over 5 minutes at five second intervals

**Supplementary movie 6D:** Video showing time lapse imaging of a hemocyte containing α-SynA53T aggregates expressing Lifeact tagged with GFP over 5 minutes at five second intervals.

**Supplementary movie 6E:** Video showing time lapse imaging of a hemocyte containing TDP 43 aggregates expressing Lifeact tagged with GFP over 5 minutes at five second intervals.

**Supplementary table 1.**

**Pixel and frame rate information**

| **Cell Type** | **Pixel width in µm** | **Frame interval in seconds** |
|---|---|---|
| WT | 0.0081 | 5.00 |
| HTTQ138 | 0.0195 | 5.00 |
| LatA treated | 0.0122 | 7.96 |
| DMSO treated | 0.0122 | 7.96 |
| Hip1 overexpression | 0.0081 | 8.82 |
| Profilin RNAi | 0.0033 | 5.00 |
| Arp2/3 overexpression | 0.0122 | 9.11 |
| Arp2/3 RNAi | 0.0037 | 5.0 |
| Hip RNAi | 0.0081 | 9.11 |
| Mrj overexpression | 0.0122 | 8.82 |

**Extended methods:**

**PIV - Drift Correction:**

Often during the process of image acquisition, there are external mechanical perturbations which lead to finite drift in the centre of mass position of the cell, with respect to the image frame, over the duration of imaging. It is therefore absolutely necessary to ensure that the internal flow vectors are corrected for drift in the cell because of the global movement. We rectified the global offset of the cell centre of mass in the time-lapse movies using StackReg, an ImageJ plugin, which can effectively correct any centre of mass movement. More details about the plugin can be found at HTTp://bigwww.epfl.ch/thevenaz/stackreg/.

Once all frames from the time-lapse movies were drift corrected, we performed the next step, namely intensity correction, which is discussed below.

**Intensity Correction**

One of the main challenges we faced while analyzing the time-lapse movies is the presence of static clathrin intensity in the background, the source of which could be clathrin patches or newly translated clathrin protein. PIV analysis of the movies in presence of this fluorescence leads to incorrect estimation of flow field. This is because the stationary fluorescent particles are considerable in number and size and show up in autocorrelation thereby significantly underestimating the flow-fields. To reduce the effect of stationary intensity during the PIV analysis, we attempted to remove the effect of stationary fluorescence in the following manner:

we first computed the mean pixel value corresponding to each pixel unit with coordinates $(i,j)$ in all the drift corrected images. This is defined as $I_{mean}(i,j) = \frac{1}{N}\sum_{l=1}^{N} I_0(i,j,l)$, where $I_0(i,j,l)$ is the intensity of a pixel $(i,j)$ in each time frame $l$ and $N$ is the number of time frames. While stationary particles will still show up in the mean image $I_{mean}(i,j)$, moving particles on the other hand will have their intensities reduced. Next, we updated each image frame using the following formula

$$I_c(i,j,k) = \max\,[0, I_0(i,j,k) - I_{mean}(i,j)] \qquad (1)$$

Here we have defined the corrected intensity $I_c(i,j,k)$ for each time-frame by subtracting the mean intensity $I_{mean}(i,j)$ from the actual intensity value. Note that we set the pixel value to 0 if mean frame intensity value is bigger. As a result of this correction, all stationary particles will disappear from each time frame, while the moving particles will have their intensities reduced as shown below.–Note that the total intensity can be written as $I_0(i,j,k) = S(i,j) + M(i,j,k)$, where $S(i,j)$ is the static intensity and $M(i,j,k)$ is the time-dependent fluorescent intensity which captures the movement of clathrin-coated vesicles. If there is no time variation in intensity within a pixel $(i,j)$, then $I_c(i,j,k) = 0$, for that pixel following Eq. (1). For cases where the time variation is non-zero, we can rewrite Eq.(1) as:

$$I_c(i,j,k) = I_0(i,j,k) - \frac{1}{N}\sum_{l=1}^{N} I_0(i,j,l) = M(i,j,k) - \frac{1}{N}\sum_{l=1}^{N} M(i,j,l). \quad (2)$$

Eq.(2) can be further re-written as,

$$I_c(i,j,k) = \frac{(N-1)}{N} M(i,j,k) - \frac{1}{N}\sum_{\substack{l=1 \\ l\neq k}}^{N} M(i,j,l) = \frac{(N-1)}{N}\left[M(i,j,k) - M_{mean}(i,j,k \neq l)\right] \quad (3)$$

Since the above equation only contains time-dependent intensity, the contribution of it to a pixel $(i,j)$ from all frames $\neq k$, will mostly be zero and as a result the quantity $M_{mean}(i,j,k \neq l)$ is expected to be a small quantity. We can thus approximate Eq.(3) as

$$I_c(i,j,k) \approx \frac{(N-1)}{N} M(i,j,k) \quad (4)$$

Due to the correction by the factor $M_{mean}(i,j,k \neq l)$, there is a decrease in the overall intensity of the moving particles. This decrease of intensity globally will not result in change in magnitudes of flow vectors during the PIV analysis. However, there will be a slight increase in the occurrence of errors while calculating the flow vectors. For this reason, we performed post-processing using standard deviation and median filters. Also note that the intensity correction method will lead to large errors if the intensity dynamics is very slow such that a given pixel is lit up for a large number of time frames, leading to a significant reduction in the estimated value $I_c(i,j,k)$. To further test the effectiveness of this algorithm, we tested this correction on a movie which does not have any static intensity. We found that while there a slight decrease in the overall intensity, the spatial distribution of the intensity profile in each time frame was invariant.

#### Supplementary Figure 1

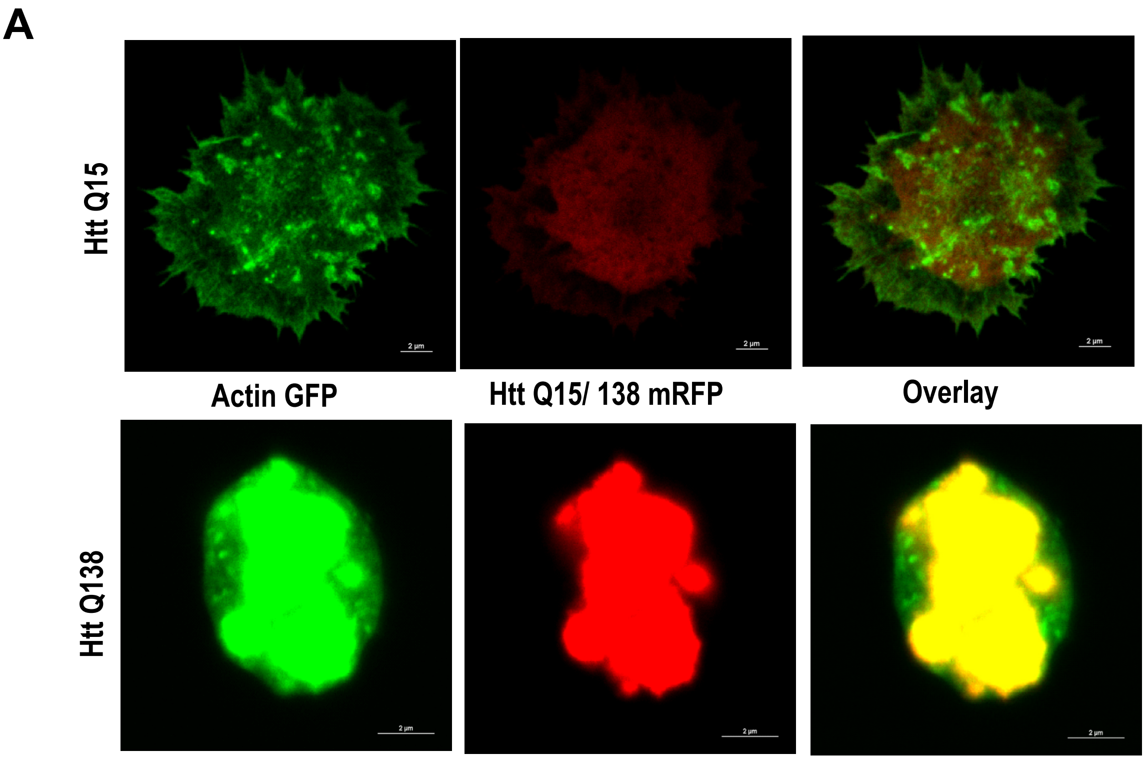


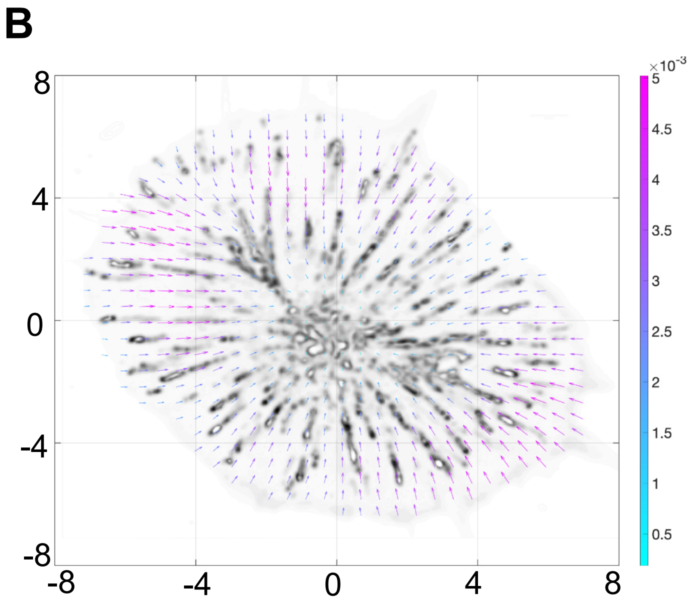

### Supplementary Figure 2

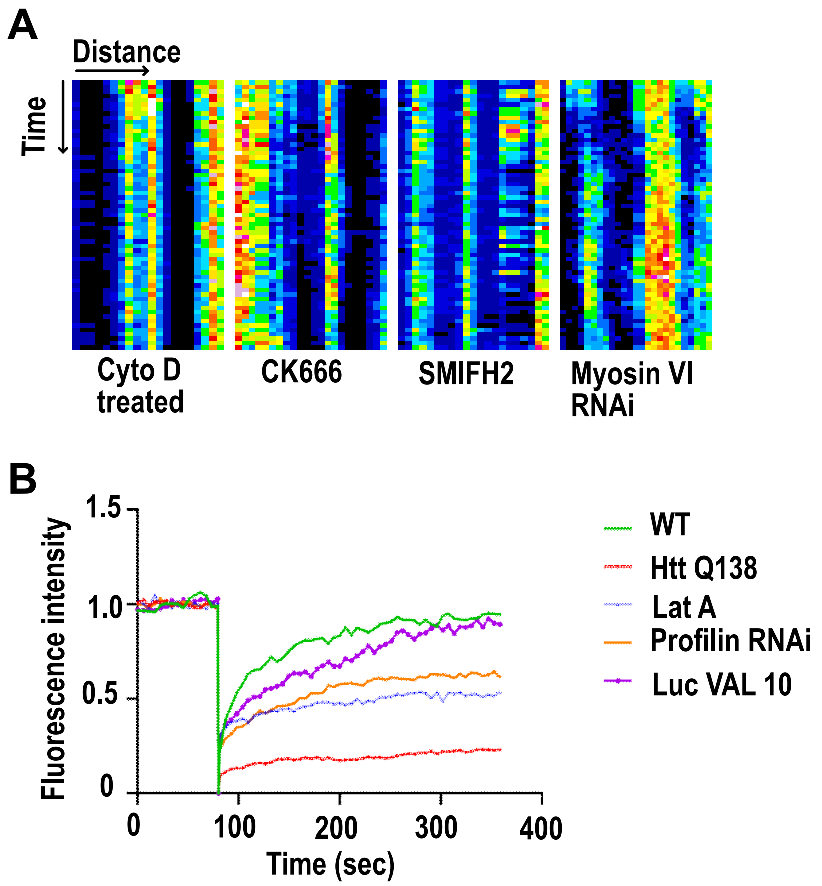

#### Supplementary Figure 3

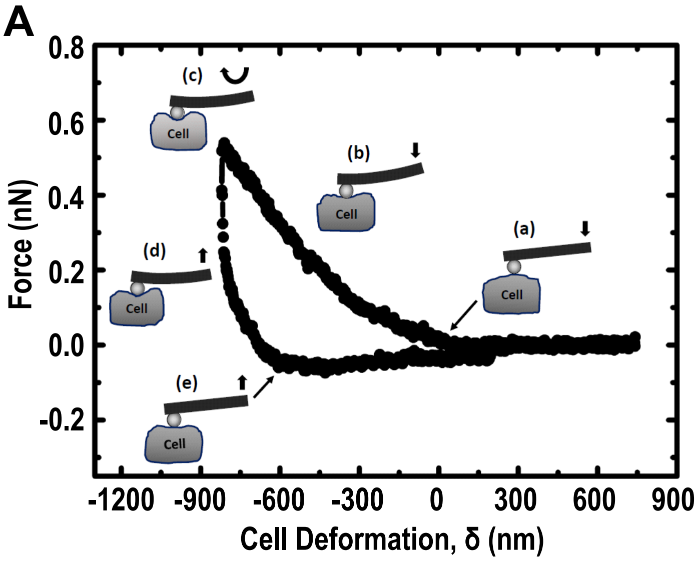

#### Supplementary Figure 4

**A**

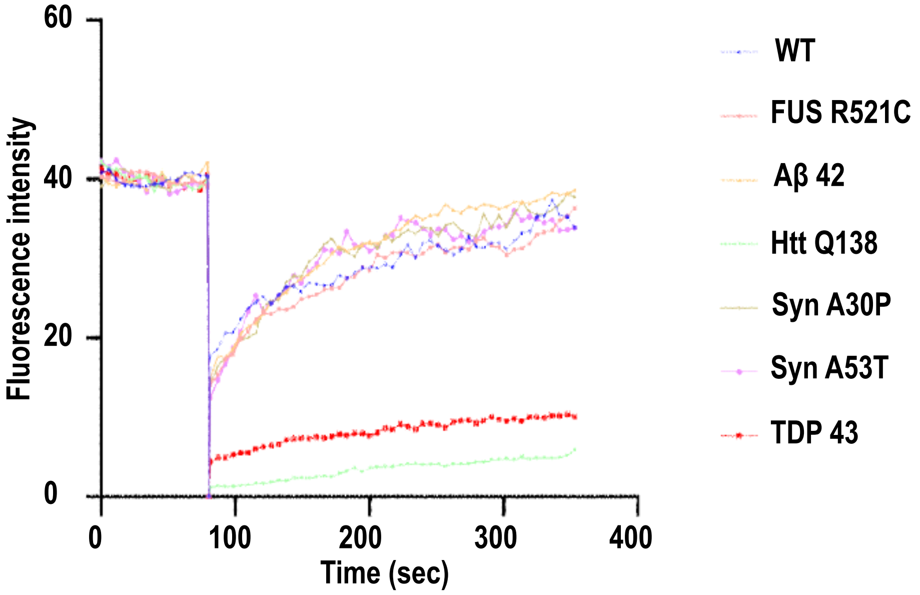

#### Supplementary Figure 5

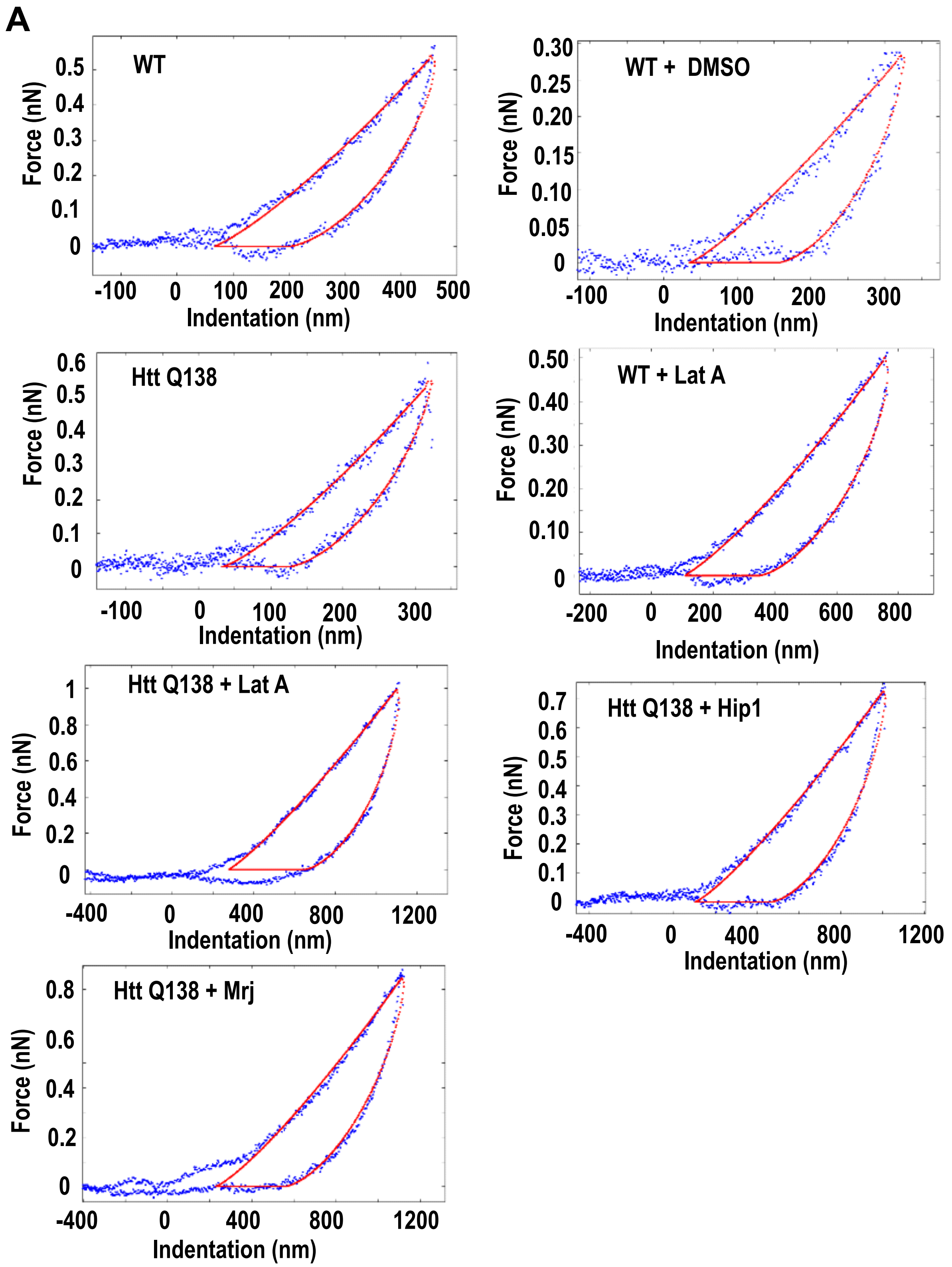
